# A dynamic foot model for predictive simulations of gait reveals causal relations between foot structure and whole body mechanics

**DOI:** 10.1101/2023.03.22.533790

**Authors:** Lars D’Hondt, Friedl De Groote, Maarten Afschrift

**Affiliations:** Department of Movement Sciences, KU Leuven, Leuven, Belgium; Department of Human Movement Sciences, Vrije Universiteit, Amsterdam, The Netherlands

## Abstract

The unique structure of the human foot is seen as a crucial adaptation for bipedalism. Its arched shape makes it possible to stiffen the foot to withstand high loads when pushing off, without compromising the range of motion. Experimental studies demonstrated that manipulating foot stiffness had considerable effects on gait. In clinical practise, altered foot structure is associated with pathological gait. Yet our understanding of how foot structure influences gait mechanics is still poor. Here we used predictive simulations to explore causal relations between foot properties and whole-body gait. Our dynamic three-segment foot model with longitudinal arch improved gait predictions compared to one- and two-segment foot models and can explain measured ankle-foot kinematics and energetics. We identified three properties of the ankle-foot complex that are crucial for healthy walking: (1) compliant Achilles tendon, (2) stiff heel pad, (3) the ability to stiffen the foot. The latter requires sufficient arch height and contributions of plantar fascia, intrinsic and extrinsic foot muscles. Insufficient foot stiffness results in walking patterns with reduced push-off power. During terminal stance plantar fascia and intrinsic foot muscles transfer energy from the metatarsophalangeal to midtarsal joint, which further increases push-off power.

## Introduction

Predictive simulations of human movement are a useful tool to explore causal relations between musculoskeletal properties and whole body gait. De novo gait patterns can be generated based on a model of the musculoskeletal system and the assumption that human locomotion is shaped by optimising performance (1,2). While simulations based on detailed 3D musculoskeletal models already capture many key features of human walking mechanics and energetics (3), predicting the ankle kinematics and kinetics remains challenging. Even the most detailed models that are currently used in predictive simulations drastically simplify the musculoskeletal system. However, insight in the effect of different simplifying assumptions on simulated gait mechanics and energetics is limited. This reflects a lack in fundamental understanding of how musculoskeletal properties shape locomotion. Here, we explored the effect of foot structure on gait mechanics. Foot models used in predictive simulations are very simple, especially in the light of the complex foot anatomy. Experimental studies suggest that foot structure has a considerable effect on walking efficiency. Yet, as it is hard to manipulate foot structure experimentally, insights gained from experimental assessments are inherently limited. Especially, our understanding of how foot structure influences foot and ankle mechanics is still poor.

Few human gait simulations were based on foot models with more than one segment but those that modelled multiple segments found a considerable effect of foot structure on gait mechanics and energetics. The foot is commonly represented by a single rigid segment (4–11), or by two segments, calcaneus-midfoot-metatarsals and toes, in predictive simulations (3,12–15). Modelling toes as a separate segment in 3D gait simulations mainly improved the accuracy of predicted knee kinematics, kinetics, and vasti activity during stance (3). Modelling a compliant foot arch as well as toes in 2D gait simulations increased the cost of transport and more so for lower speed (16). Arch stiffness influenced the cost of transport (17). However, the effect of modelling the foot arch on gait kinematics and kinetics was not reported. These prior studies demonstrate the importance of the foot for whole body gait mechanics and energetics, and motivate a more profound investigation of foot models in 3D gait simulations. Here we modelled the different foot structures and properties that influence foot stiffness: the longitudinal arch, soft tissues contributing to foot stiffness, and muscles actuating the foot.

The compliant longitudinal foot arch has been suggested to improve gait efficiency by storing and releasing elastic energy (18). The foot absorbs energy while it is flat on the ground while the body centre of mass moves forward over the foot. Energy is mainly stored by deformation of the medial longitudinal arch, stretching of long and short plantar ligaments, and plantar fascia (18–20). Energy release at push-off is believed to enable efficient generation of high push-off power (18–20). Although elastic energy is stored in the foot during both walking and running, this energy storage seems to contribute more to running than to walking efficiency (20). Modelling a compliant longitudinal foot arch will provide more insight in its contribution to energy storage and release in the foot.

During walking, an energy efficient push-off depends on the ability of the foot to act as a stiff lever. Most push-off power comes from the triceps surae and Achilles tendon (21). While the body moves over the standing foot, energy is stored in the Achilles tendon. Releasing the stored energy generates a powerful forefoot push-off (21,22). During push-off, the foot acts as a lever between the Achilles tendon and the ground (23). A rigid foot is essential to ensure an efficient push-off without energy dissipation in the foot (19,24). It has been suggested that the windlass mechanics stiffens the foot at push-off. In 1954, Hicks described how extending the toes tensions the plantar fascia by winding it around the metatarsal heads (24). This then pulls the base of the medial longitudinal arch together, raising the arch and stiffening the foot (24). To be effective, the windlass mechanism requires the plantar fascia to be very stiff, which would hinder its ability to store elastic energy. A model including plantar fascia will allow us to study the contribution of the windlass mechanism to stiffening the foot at push-off.

Intrinsic foot muscles can modulate foot stiffness and might therefore play an important role in optimizing efficiency by enabling the foot to function both as an elastic structure that stores and releases energy, and as a rigid lever. The intrinsic muscles in the superficial layers of the plantar aspect of the foot have short fibres, a high pennation angle and a long tendon (25,26). Muscles with similar architecture are suited to produce force to modulate the stiffness of their tendon and to absorb and return elastic energy in a controlled way (27). Multiple plantar intrinsic muscles originate on the calcaneus and insert on different toe bones (25,26). Contraction of these muscles will increase the apparent stiffness of the plantar fascia, and the foot as a whole (28). Electromyographic recordings showed that intrinsic foot muscles are active during the stance phase of walking (28–30). Activity is highest in the second half of the stance phase, when the triceps surae contract and the ankle plantarflexion moment is high (28,29). Preventing intrinsic muscle contraction by injecting a nerve block decreased push-off power of the ankle-foot complex, yet arch deformation and cost of transport were unaffected (31). Modelling intrinsic foot muscles will allow us to elicit their contribution to foot stiffness and walking mechanics.

Gait efficiency might depend on well-tuned interactions between foot and ankle structures. Foot stiffness influences the ability for rapid energy release from the Achilles tendon. Therefore, we also investigated how Achilles tendon stiffness influenced simulated gait mechanics. Furthermore, foot stiffness does not only depend on arch stiffness but also on the mechanical properties of the soft tissues between the bones and foot sole. Especially at initial contact when only the heel is in contact with the ground, soft tissues might be the main contributor to foot stiffness. The visco-elastic fat pad under the heel has been shown to absorb the shock and decelerate the lower leg at initial contact (22,32). Therefore, we also explored the stiffness of the foot-ground contact.

Here we present a dynamic three-segment foot model (hindfoot, midfoot-forefoot, and toes) for 3D predictive simulations of gait. The model captures passive stiffness of the midtarsal joint and includes the plantar fascia. The foot is actuated by extrinsic as well as intrinsic foot muscles. The presented foot model improves gait predictions compared to one- and two-segment foot models (3,6) by better capturing ankle, midtarsal, and metatarsophalangeal kinematics as well as soleus and peroneus longus activity. We evaluated the effect of modelling choices on the predicted gait mechanics and energetics. We identified a combination of model parameters that are important to generate physiologically plausible gait kinematics: (1) Achilles tendon stiffness, (2) contact stiffness, (3) foot arch height, (4) plantar fascia stiffness, and (5) muscle actuation of the foot arch.

## Material and methods

### Musculoskeletal model

We adapted the musculoskeletal model used by Falisse et al. (3) – which is based on OpenSim’s gait2392 model (33,34) and the model proposed by Hamner et al. (35) – to include a midtarsal joint, plantar fascia, and a plantar intrinsic foot muscle. The resulting model has 33 skeletal degrees of freedom (dofs) (pelvis as floating base: 6 dofs, hip: 3 dofs, knee: 1 dof, ankle: 1 dof, subtalar: 1 dof, midtarsal: 1 dof, metatarsophalangeal (MTP): 1 dof, lumbar: 3 dofs, shoulder: 3 dofs, and elbow: 1 dof). We used Newtonian rigid body dynamics to model skeletal motion (36,37). The lower limb and lumbar joints are actuated by 94 Hill-type muscletendon units (92 muscles according to the gait2392 model and the right and left plantar intrinsic foot muscles as described below) (38,39). Muscle excitation-activation coupling is given by Raasch’s model (40). The shoulder and elbow joints are actuated by ideal torque actuators (3). Each joint has viscous friction (coefficient 0.1 Nm s rad^-1^) and exponential stiffness (13) to represent the lumped effects of unmodelled soft tissue (6).

A first series of model adaptations aimed at better capturing triceps surae properties. Muscle parameters of the generic model are based on elderly cadavers (41). Age related muscle atrophy results in underestimations of maximal isometric forces (42,43). On the other hand, a model with maximal isometric forces based on in vivo imaging of young adults, which are about twice as large (43), overestimated passive joint moments, maximum isometric joint moments, and passive force contributions during walking (42). Therefore, we opted for a moderate increase of 20% in the maximal isometric force of the triceps surae. Based on a sensitivity analysis (supplement S1) and previous work (3,44), we reduced the Achilles tendon stiffness to half its default value (38). We adapted the parameters related to passive stiffness of the muscles spanning the ankle joint as the current passive ankle moment-angle relation does not match experimental data (45). Passive stiffness of all muscles crossing the ankle joint was increased by shifting their passive force-length characteristic (38) with 10% of the optimal fibre length towards smaller fibre lengths. The resulting passive ankle moments are consistent with measurements (45) (Figure S2). We calculated peak isometric ankle moments by assuming maximal activation of agonists and zero activity in antagonists. The model-based ankle moment is lower than the data presented by Holzer et al. (46), but consistent with the data presented by Anderson et al. (47), Marsh et al. (48), and Sale et al. (49) (Figure S4). We evaluated how each of these changed muscle parameters influence the predicted gait (Figure S1,S3,S5).

We changed the orientation of the MTP joint axis to obtain more realistic behaviour of the flexor digitorum longus tendon. Assuming an MTP axis normal to the sagittal plane causes the moment arm of the flexor digitorum longus with respect to the metatarsophalangeal joint to decrease with increasing MTP extension. For extension of 30° and more, the moment arm is lower than 3 mm. This is inconsistent with the anatomy of the tendons passing over the metatarsal heads. We avoided this by modelling an MTP axis oriented according to the MTP plantarflexion/dorsiflexion axis proposed by Malaquias et al. (50).

We simplified the foot arch by modelling it as a midtarsal joint with a single rotational degree of freedom, connecting calcaneus and midfoot segments. Midfoot and forefoot segments are rigidly connected. Midtarsal joint centre, segment definitions, and segment mass properties were taken from Malaquias et al. (50). We simulated walking for models with six different midtarsal joint axis orientations. Orientations 1 and 5 are respectively the anterior-posterior and oblique axis defined by Malaquias et al. (50). Orientations 2, 3, and 4 are interpolations at 30° increments (supplement S4). The sixth axis orientation is normal to the sagittal plane. Walking simulations with axis orientation 4 resulted in the lowest optimal cost, thus we selected this orientation for the final model (Figure 1). We computed the rotational stiffness of the midtarsal joint by combining the contributions of individual ligaments. We included long plantar ligaments, calcaneo-navicular (plantar and bifurcate) ligaments, and calcaneo-cuboid (plantar, dorsal, and bifurcate) ligaments as modelled by Malaquias et al. (50). We used a stiffening stress-strain characteristic (51) but replaced the polynomial function by an exponential function that closely fitted the original polynomial function to improve computational stability.

**Figure 1.**
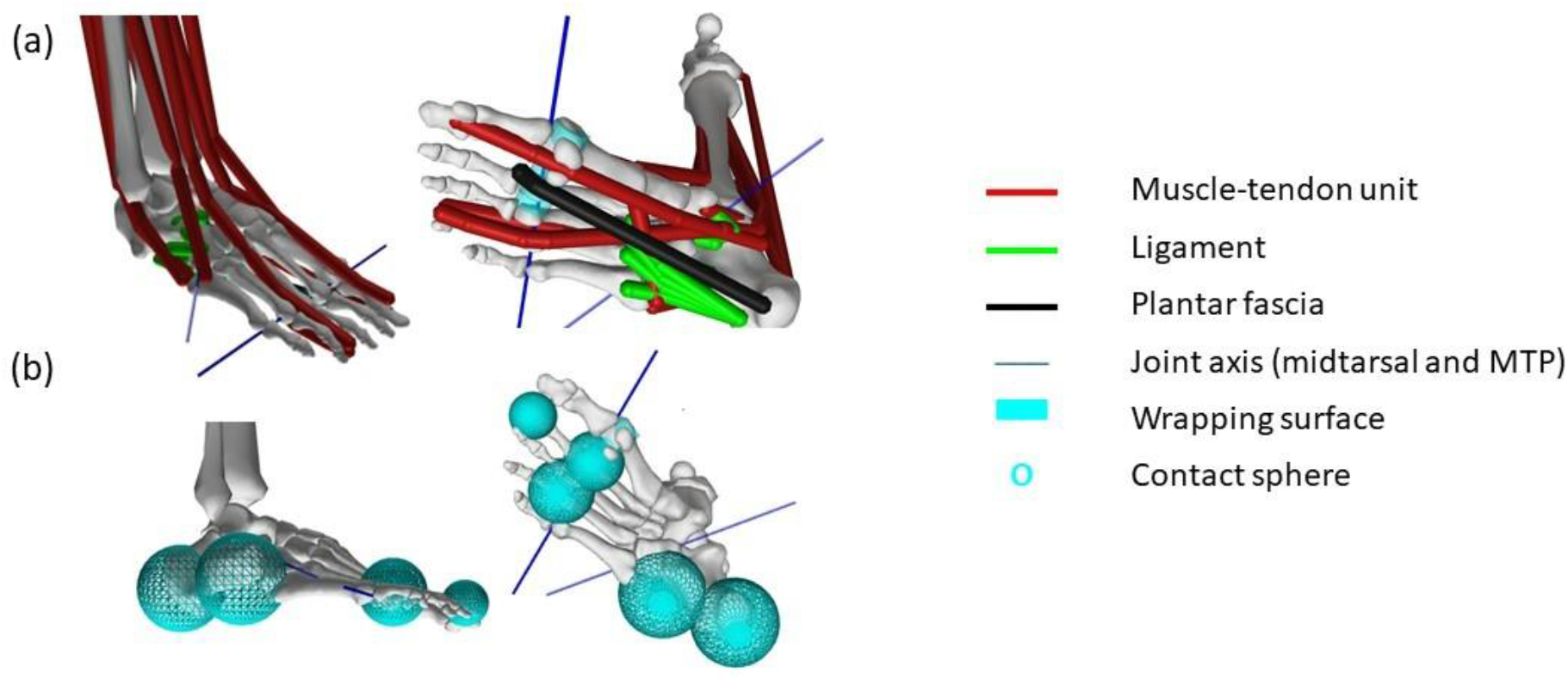
(a) Musculoskeletal geometry of the ankle and foot. Plantar intrinsic muscle is not visible, because it shares the same path as the plantar fascia. (b) Location and size of contact spheres: two spheres are attached to the calcaneus, two to the forefoot, and one to the toes.

The plantar fascia is modelled as a single elastic element with origin on the calcaneus and insertion on the second proximal phalange (50), which wraps around a cylinder with radius 9.5 mm at the metatarsal head (52,53). Based on cadaveric data (54), we set a nominal plantar fascia cross-section of 60 mm^2^. We assumed a purely elastic plantar fascia, and compared different stress-strain characteristics. We simulated walking with the plantar fascia models presented by Natali et al. (55) (we refer to this model as “Natali2010”) and Gefen (51) (“Gefen2002”). Parameters of both models were identified from uniaxial tension tests of plantar fascia samples (51,55). As both models are based on experimental data there is no clear criteria for selecting one over the other. We used the model from Natali et al. as this resulted in lowest optimal cost in walking simulations. Additionally, we compared gait simulations with these two models to simulations with a plantar fascia with Young’s modulus 350 MPa (56), and the model used by Song et al. (16) (Figure S11). Plantar fascia slack length (146 mm) was chosen such that a standing foot, bearing only the weight of foot and tibia, is in anatomical position (i.e. midtarsal and MTP joint angles are zero).

We modelled the plantar intrinsic muscles of a foot as a single Hill-type muscle, because they act as a functional unit (30). Origin, insertion, and wrapping of the muscle are the same as the plantar fascia. Parameters describing the muscle are derived from cadaveric measurements of flexor digitorum brevis (FDB) and abductor hallucis (AH) (25). Based on the reported muscle belly of 73 mm and ratio of fibre length over muscle length of 0.27 (25), average fibre length of FDB is 19.7 mm. Maximal contraction velocity is 10 optimal fibre lengths per second (6). Pennation angle at optimal fibre length is 20°, equal to measurements of FDB and AH (25). To determine the physiological cross sectional area (PCSA) of the lumped muscle, we sum the PCSA of FDB and AH for a total of 500 mm^2^ (25). Total PCSA is consistent with results from ultrasound measurements in healthy young adults (57). We double the PCSA, to account for the contributions of other unmodeled plantar intrinsic muscles that cross both midtarsal joint and MTP joint: (1) abductor digiti minimi originates on the calcaneus and inserts on the fifth proximal phalanx (26), (2) quadratus plantae originates on the calcaneus or the long plantar ligament and inserts on the flexor digitorum longus tendon (58), (3) flexor hallucis brevis originates on the tendon of the tibialis posterior and inserts on the first proximal phalanx (59). In our model, we do not consider the specific anatomy of these three muscles. Assuming a specific tension of 70 N cm^-2^ (6), the max isometric force is 700 N. To model the stiffness provided by the internal tendon in highly pennate muscles, we shift the passive force-length characteristic (38) with 10% of the optimal fibre length towards shorter fibre lengths.

We determined tendon slack length based on the assumption that muscle fibres work close to their optimal length during walking. Since we modelled the muscle-tendon unit along the same path as the plantar fascia, it has the same length as the plantar fascia. The plantar fascia is expected to have 0 – 4% strain while walking (56,60) allowing us to estimate the expected muscle-tendon length based on the selected plantar fascia slack length. We then calculated normalized fibre lengths for different tendon slack lengths assuming an isometric contraction at an activation of 0.4. We selected the tendon slack length (125 mm) that resulted in fibre lengths around the optimal fibre length. To test the influence of parameter choices, we repeated the walking simulations with different values for optimal fibre length, tendon slack length, and maximal isometric force (supplement S5). We also tested a model without plantar intrinsic muscles.

The model was scaled to the anthropometry of the subject (see below) with the OpenSim scale tool (33). Scale factors were calculated based on marker data of the static trial. For the foot (talus and distal segments), we included manual measurements to determine non-uniform scale factors. Anteroposterior scaling is based on the distance between the dorsal calcaneus marker and the hallux marker. Mediolateral scaling is based on the distance between the markers on the heads of the first and fifth metatarsal. In the vertical direction, the vertical distance between centres of medial malleolus and first metatarsal head (based on manual measurements) determines the size of the foot. Because scaling of the foot is non-uniform, insertion points of tibialis anterior and posterior, and peroneus (brevis, longus, and tertius) were distorted. We substituted these insertion point locations from an alternative musculoskeletal model (50), after scaling it to match our subject-specific model. Foot proportions of this alternative model were close to the subject’s foot proportions, thus scaling did not cause distortions. Optimal fibre length and tendon slack length of the adjusted muscles were linearly scaled (cfr. the OpenSim scaling method for these parameters (33)). To test the effects of this scaling method, we also created a model with uniformly scaled feet. The uniform scale factor is proportional to the distance between the dorsal calcaneus marker and the hallux marker. Uniform scaling results in a lower foot arch compared to non-uniform scaling (Figure 9e).

We used smoothed Hunt-Crossley contact spheres (61,62) to model foot-ground contact. Each foot has five contact spheres: heel pad, lateral longitudinal arch, ball of the foot (2 spheres), and toes. The location and radius of each sphere were chosen to represent areas with high plantar pressure during standing and walking (54,63–65), and based on anatomical reference (66). To tune the vertical position of the contact spheres, we tracked experimental joint angles from the static trial with skeletal and contact dynamics. We solved the static force equilibrium, and adjusted the vertical contact positions to reduce offset with respect to experimental pelvis-to-ground coordinates. We set the stiffness parameter of the contact model (i.e. plain strain modulus) to 10 MPa, instead of the 1 MPa used previously (3,6). Preliminary simulations showed that this improved the realism of the contact deformation during walking. We evaluated how configuration and stiffness of the contact spheres influence the predicted gait (supplement S12).

We also present an alternative 2-segment model. It has the same complexity as the state-of-the-art foot model (3), with parameter values based on our novel 3-segment model. In the 2-segment model, the midtarsal joint is locked in anatomical position. The toes are not actuated by muscles. Instead, the passive MTP joint has a spring (20 Nm/rad) and damper (2 Nm s/rad) (3).

### Simulations of standing foot under vertical loading

We replicated a series of experiments that evaluated the deformation of the foot under external loading (18,67,68) in simulation and compared simulated and experimental outcomes to validate our model. Foot stiffness is commonly assessed by applying vertical forces to a standing foot and measuring the deformation (18,51,56,67–69). To simulate such experiment, we considered only the right tibia and foot of our musculoskeletal model. In these simulations, the tibia had 3 degrees of freedom with respect to the ground while translations in the transverse plane and rotation around the vertical axis were locked. The angle between toes and ground was fixed (cfr. (67,68)), and muscle activity was set to zero for cadaver experiments and to a small baseline activation (0.01) for in vivo experiments. We imposed a vertical force on the tibia and solved for static equilibrium. We repeated this for different force values within the experimental range to obtain force-deformation curves.

First, we simulated an experiment from Ker et al. that assessed longitudinal elongation of the foot under vertical loading in different conditions by subsequently removing more soft tissue (18). To replicate this experiment, we used a model in which individual ligaments were represented (before lumping them for computational efficiency in gait simulations). Ker et al. did not discuss if and when intrinsic muscles were cut. We considered removing intrinsic muscles as an additional step after removing plantar fascia, and before long plantar ligament. To compare simulation outcomes to experimental data, which was obtained on the foot of an 85 kg male, we scaled forces by the ratio of body weights (0.76) and elongation by the square root of this ratio (0.87).

Next, we simulated two experiments that evaluated the effect of flexing or extending the toes on arch compression. Yawar et al. tested how extending the toes (15°) influenced the vertical compression of a cadaver foot (defined as vertical displacement of the transected tibia), compared to having the toes in neutral position (68). Welte et al. applied a vertical load up to body weight on the thigh of a seated person (force was aligned to be vertical to navicular) and assessed how flexing or extending the toes by 30° influenced arch compression (67). Arch compression was defined as the change in vertical position of a marker on the navicular normalised to the maximal value for the subject (67).

### Simulations of walking

We used our previously developed framework (6,36) to simulate walking. We formulated gait as an optimal control problem. We solved for muscle excitations that minimise an objective function, constraint by the musculoskeletal dynamics and prescribed average forward velocity of the pelvis (1.33 m s^-1^, preferred walking speed of the subject). We imposed symmetry between left and right steps, which limits the time horizon to half a gait cycle. The objective function is a weighted sum of metabolic energy squared, the sum of squared muscle activations, the sum of squared joint accelerations, and the sum of squared joint limit torques, integrated over the time horizon and divided by the distance travelled. We calculated metabolic energy of muscles with the model presented by Bhargava et al. (70), which we made continuously differentiable by approximating conditional statements with a hyperbolic tangent (supplement S6). This framework is implemented in MATLAB (The Mathworks Inc., USA). We used direct collocation with a third order Radau polynomial basis. We used algorithmic differentiation (CasADi 3.5.5 (71)) to calculate the derivatives required for gradient based optimisation. The optimisation problems were solved with IPOPT (72) and MUMPS (73). For a detailed description of the simulation framework, we refer to (6).

Based on convergence analysis we selected 100 mesh intervals, cold-start initial guess (Figure S15) and convergence tolerance of 1e-04 (Table S2). All results shown are obtained from simulations with these settings.

To evaluate the sensitivity of predicted gait to differences in the musculoskeletal model, we performed simulations for a range of models and model parameters.

### Experimental data

We compared simulation outcomes to previously published experimental data (3). The data (marker trajectories, ground reaction forces, surface electromyography) was collected from a healthy female adult (mass: 62 kg, height: 1.70 m, age: 35) during 10 gait cycles of overground walking at self-selected speed (1.33 m/s) (3). The subject was instrumented with 59 retro-reflective skin-mounted markers. There were 10 markers mounted on each foot (supplement S8). Additionally, we measured (with measuring tape) the normal distance from the medial malleolus centre and the first metatarsal head centre to the ground while the subject was in a seated position with the foot flat on the ground. We scaled a musculoskeletal model to the anthropometry of the subject, as described in the section on the musculoskeletal model. We performed inverse kinematic and inverse dynamic analysis (34) based on the model with 3-segment feet and selected midtarsal joint orientation (Figure 1), and calculated joint powers. We repeated inverse kinematic analysis for models with different midtarsal joint axis orientations (Figure S7).

## Results

### Simulations of standing foot under vertical loading to validate model parameters

To evaluate the realism of our foot model, we replicated a series of experiments that applied controlled loads to the foot in simulation.

Loading the proposed foot model by a vertical force up to 3 kN resulted in a maximal longitudinal elongation of 4.2 mm (Figure 2a), which is lower than the 6 to 8 mm observed by Ker et al. for an intact cadaver foot (18). When using the plantar fascia stress-strain characteristic proposed by Gefen et al. instead of that proposed by Natali et al., the simulated elongation was 6.1 mm and thus in closer agreement with experimental data. The shape of the force-elongation curve was in agreement with experimental data. Overall, our model captured the contribution of the different ligaments and passive muscle forces to foot stiffness as can be seen by the agreement between simulated and experimental force-elongation curves after sequentially removing soft tissues (Figure 2a). After removing all ligaments, the remaining stiffness of the model comes from coordinate limit torques, which we included to represent unmodelled soft tissue contributions (6).

**Figure 2:**
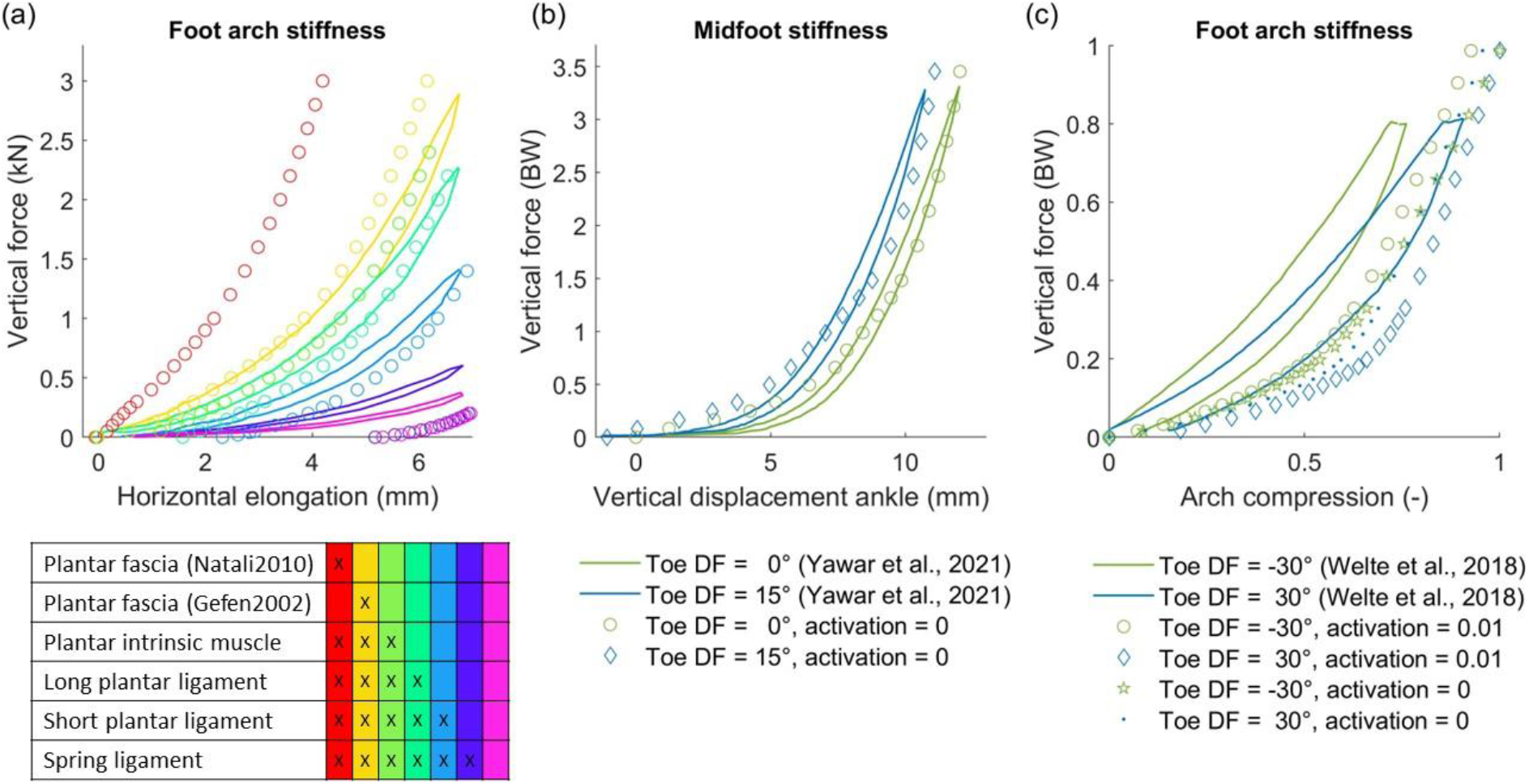
Deformation of a standing foot under vertical compression. Full lines indicate experimental data (loading and unloading curve), circles represent simulations. (a) Repeated loading of a cadaver foot after subsequently removing more soft tissue. Reference data was digitised from Ker et al. (18). The table gives on overview of structures that are present in each condition. (b) Vertical force-displacement curve when the toes are in anatomical position, and when the toes are in 15° dorsiflexion (DF). Reference data was digitised from Yawar et al. (68). We imposed muscle activation = 0 to represent a cadaver. (c) Vertical loading of a standing foot of an in vivo subject, with toes passively dorsiflexed (DF) 30° and −30°. Reference data was digitised from Welte et al. (67). We assume baseline muscle activation (0.01) for the in vivo subject. To test whether the experimental outcome would change in a cadaver, we repeated the simulations without muscle activation.

Similar as in experiments we found that imposing a different toe position altered the relation between the applied vertical load and vertical compression of the foot. In agreement with experimental data obtained from a cadaver foot, extending the toes 15° shifted the force-compression curve towards lower displacements, but did not affect the shape of the curve (i.e. foot stiffness remains constant). We predicted a 1.2 mm shift (Figure 2b), compared to 1.3 mm observed in experiments (68). In vivo, dorsiflexing the toes (30°) reduced the stiffness of the foot arch (i.e. reduced the slope of the force-compression curve) relative to having plantarflexed toes (30°) (Figure 2c) (67). Our simulations capture the change in stiffness with toe flexion when we impose a small baseline activity to the muscles (0.01) - consistent with people never fully relaxing their muscles - but not in the absence of any activity. Our simulations slightly overestimate compression with respect to Welte et al., especially for smaller loads. Note that our model does not capture the observed hysteresis throughout all experiments, because we assumed static equilibrium at every discrete load.

### Gait simulation with a 3-segment foot model

We compared simulated gait patterns using the proposed 3-segment foot model to experimental data to evaluate the realism of our simulations, as well as to simulations based on the 2-segment model we used before (Falisse et al. (3)) (Figure 3, black and orange traces).

**Figure 3:**
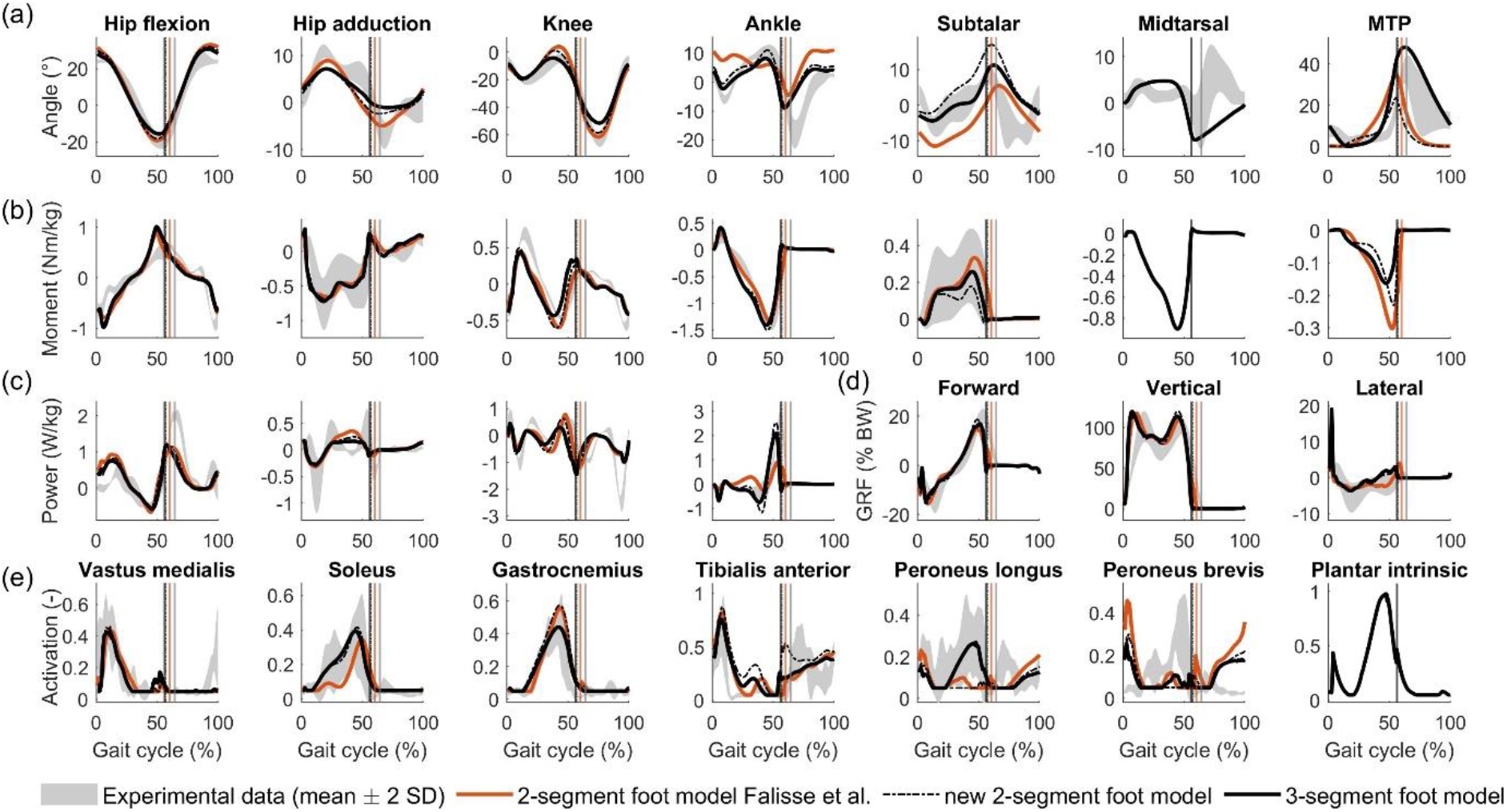
predictive simulation results with the presented model compared to experimental data and state-of-the-art simulation. Vertical lines indicate stance-to-swing transition. (a) Kinematics. (b) Kinetics. (c) Joint powers. (d) Ground reaction forces (GRF), expressed as % body weight (BW). (e) Muscle activation. Gastrocnemius indicates the medial gastrocnemius. Kinematics and kinetics of all joints can be found in supplement S9.

Using the proposed 3-segment foot model improved the agreement between simulated and experimental ankle and foot kinematics compared to the 2-segment foot model from Falisse et al. (3). Whereas simulations based on the 3-segment foot model capture experimental ankle kinematics during the stance better than simulations based on the 2-segment foot model, they still underestimate plantarflexion at the end of push-off (Figure 3). Using our 3-segment foot model also clearly improved the prediction of subtalar, midtarsal and MTP angles compared to using Falisse’s 2-segment foot model (Figure 3). Using our 3-segment foot model also led to slightly different estimates of hip and knee kinematics than using Falisse’s 2-segment foot model.

Simulations based on our 3-segment foot model resulted in similar ankle, knee and hip moments and ground reaction forces as simulations based on Falisse’s 2-segment foot model. The main differences are a slight underestimation of the knee flexion moment during push-off and a sight overestimated of the ankle plantarflexion moment during the stance phase when using our 3-segment model as compared to Falisse’s 2-segment model. We could not evaluate the realism of the predicted midtarsal and MTP moments as we did not measure ground reaction forces acting on each foot segment. Simulated ground reaction forces are similar for both foot models. Simulations based on both models predict the M-shape pattern but display a steeper increase after initial contact compared to the measured ground reaction forces.

Using our 3-segment model instead of Falisse’s 2-segment model improved the prediction of the ankle joint power. The peak ankle power obtained with the 3-segment foot model (~ 2 W/kg) is similar to the experimental peak ankle power (~ 2.5 W/kg) and is clearly a better prediction than the peak power obtained with Falisse’s 2-segment foot model (~ 1 W/kg). Simulated hip and knee powers are similar for the 3-segment and 2-segment foot models. Both models capture the pattern of measured knee and hip power but underestimate peak powers.

Using our 3-segment model instead of Falisse’s 2-segment model also improved agreement between simulated and experimentally observed activity of muscles actuating the ankle and foot joints. In contrast to the 2-segment foot model, simulation with the 3-segment foot model capture the early onset and slow build-up of soleus activity as well as peroneus longus activity. Yet simulated peroneus brevis activity remains inaccurate in both models. Consistent with published measurements of flexor digitorum brevis and abductor hallucis, the plantar intrinsic muscle is most active during the second half of stance (28,30) with a peak shortly before toe-off and a small peak after initial contact (29).

Simulated foot and ankle energetics are mostly in agreement with experimental results obtained from unified deformable segment analysis reported in literature (for details, see supplement S10). Simulated power curves for different parts of the foot (i.e. distal to hallux, forefoot, hindfoot, and shank) capture the results obtained by Takahashi et al. (74), with exception of two features (Figure 4a). First, our simulations largely overestimate the negative power distal to shank and hindfoot after initial contact. This peak is attributed to absorption of impact energy after heel strike (75). Overestimating impact is consistent with the overestimation of the rise in ground reaction forces after initial contact (Figure 3d). Second, we underestimate negative power distal to the hindfoot, and overestimate negative power distal to the forefoot during terminal stance (Figure 4a). Our simulation has less negative and net work distal to the hindfoot compared to experiments, but captures the observation that the net work distal to the shank is low (Figure 4b), i.e. the ankle-foot complex is close to energy-neutral during walking (74).

**Figure 4:**
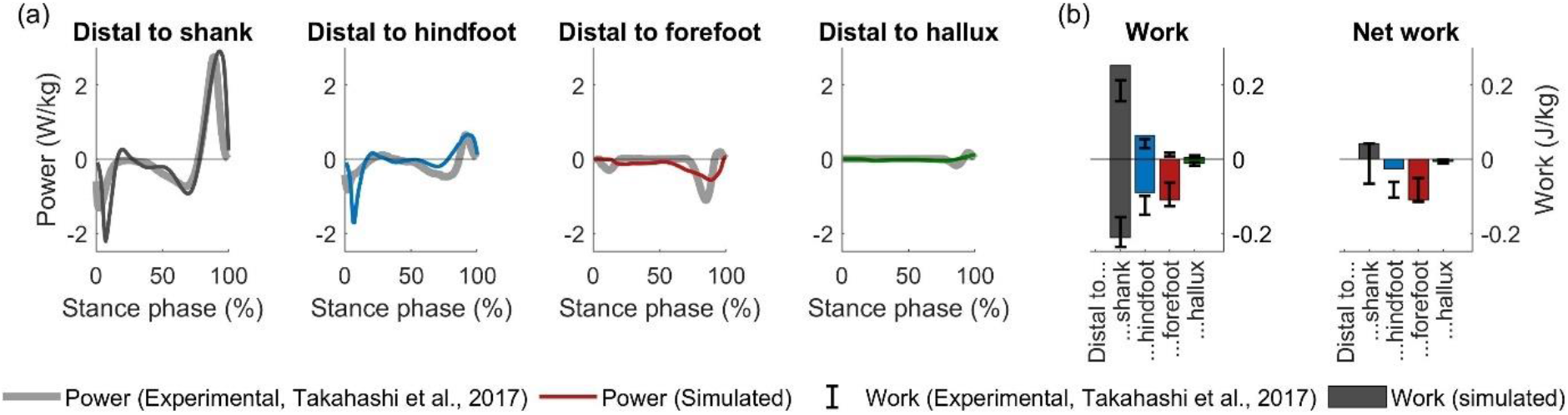
Deformation power and work by soft tissue distal to ankle-foot segments. (a) Power distal to segments. (b) Positive, negative, and net work. Experiment-based data, published by Takahashi et al. (74), is plotted as reference.

### Effects of modelling choices

We evaluated the effect of modelling choices on the predicted walking motion to explore how different foot structures shape walking mechanics and energetics. In each of the following paragraphs we compare the effect of a specific modelling choice with respect to the nominal model. We discuss only the gait features that are sensitive to the change. For additional outcomes, we refer to the supplementary material.

### Number of foot segments

Our 3-segment model and Falisse’s 2-segment model not only differ in the number of segments but also in maximal isometric force of the triceps surae, stiffness of the Achilles tendon, passive stiffness of the muscles crossing the ankle, height of the foot, and stiffness and geometry of the foot-ground contact. To dissociate the effect of modelling the foot in more detail versus altering stiffness parameters, we changed all these parameters in Falisse’s 2-segment model to match the values in our 3-segment model. From now on, we will refer to this model as the nominal 2-segment model. Simulations based on the nominal 2-segment model capture the experimentally observed ankle dorsiflexion and ankle power during the stance phase (Figure 2 dotted lines, supplement S6). Yet the simulations based on the nominal 2-segment foot model fail to capture fine details of foot kinematics.

### Foot-ground contact stiffness

Foot-ground contact stiffness had a large influence on knee flexion and the knee extension torque during early stance when using the 3-segment foot model but not when using the nominal 2-segment model. More compliant foot-ground contact (stiffness 1 MPa instead of 10 MPa) resulted in increased stance knee extension and reduced stance knee extension moment in the 3-segment model (Figure 5). A model with stiff contact results in a simulated peak heel pad compression of 7.7 mm, which is still slightly higher than the 3.8 – 4.8 mm reported by Gefen et al. (76). Further increasing contact stiffness reduces compression, but does otherwise not affect the predicted gait (Figure S20).

**Figure 5:**
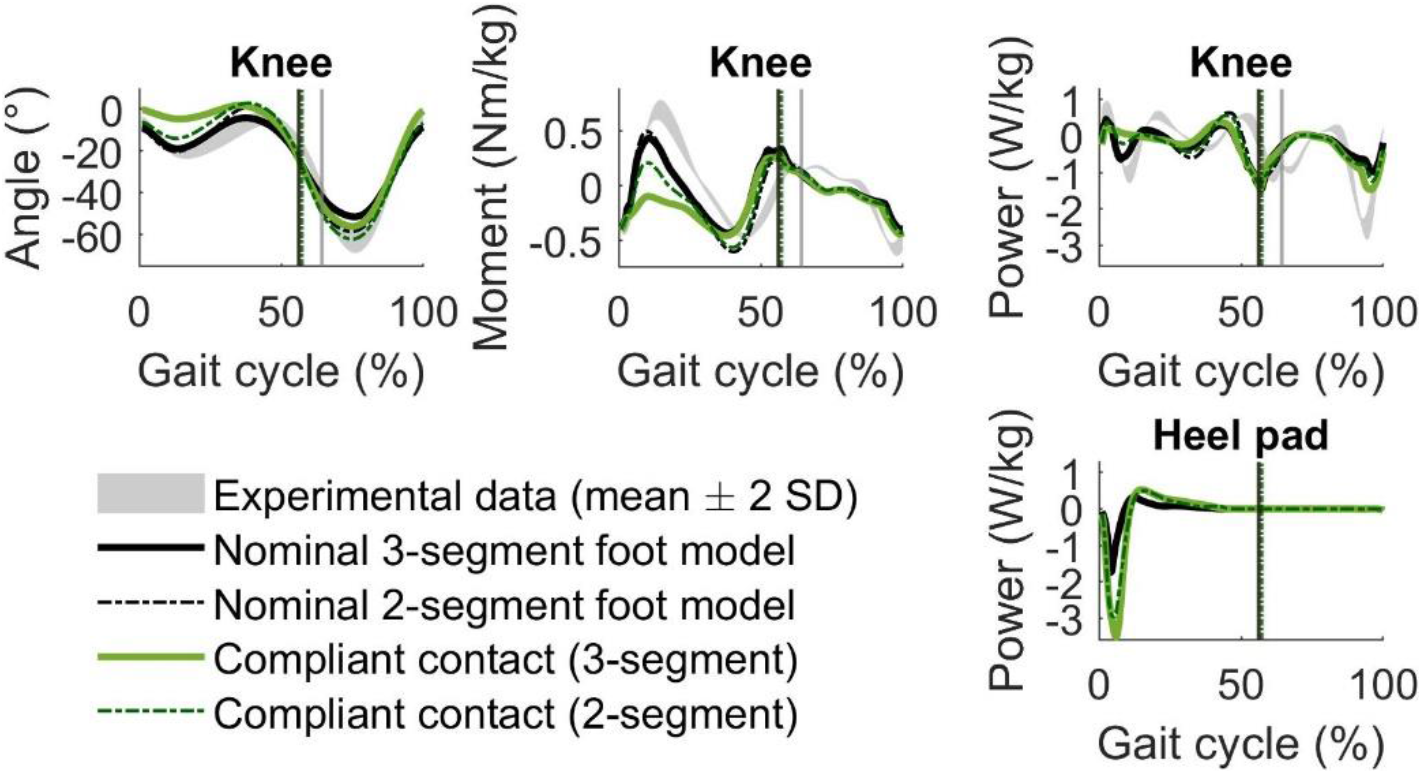
Effect of foot-ground contact stiffness. Reduced stiffness of the foot-ground contact influenced knee mechanics and heel pad deformation power.

### Achilles tendon stiffness

We found that a sufficiently compliant Achilles tendon was important to predict ankle dorsiflexion during mid-stance. In both our 3-segment and nominal 2-segment foot models, high Achilles tendon stiffness (i.e. generic normalised tendon stiffness (38)) results in little or no ankle dorsiflexion during midstance in contrast to experimental observations (Figure 6). The energy storage in the Achilles tendon and ankle push-off power are lower in the simulations with a stiff Achilles tendon than in the simulations with the lower default tendon stiffness that we propose here. Using a higher versus lower Achilles tendon stiffness also leads to a worse agreement between simulated and experimental soleus activity (Figure 6).

**Figure 6:**
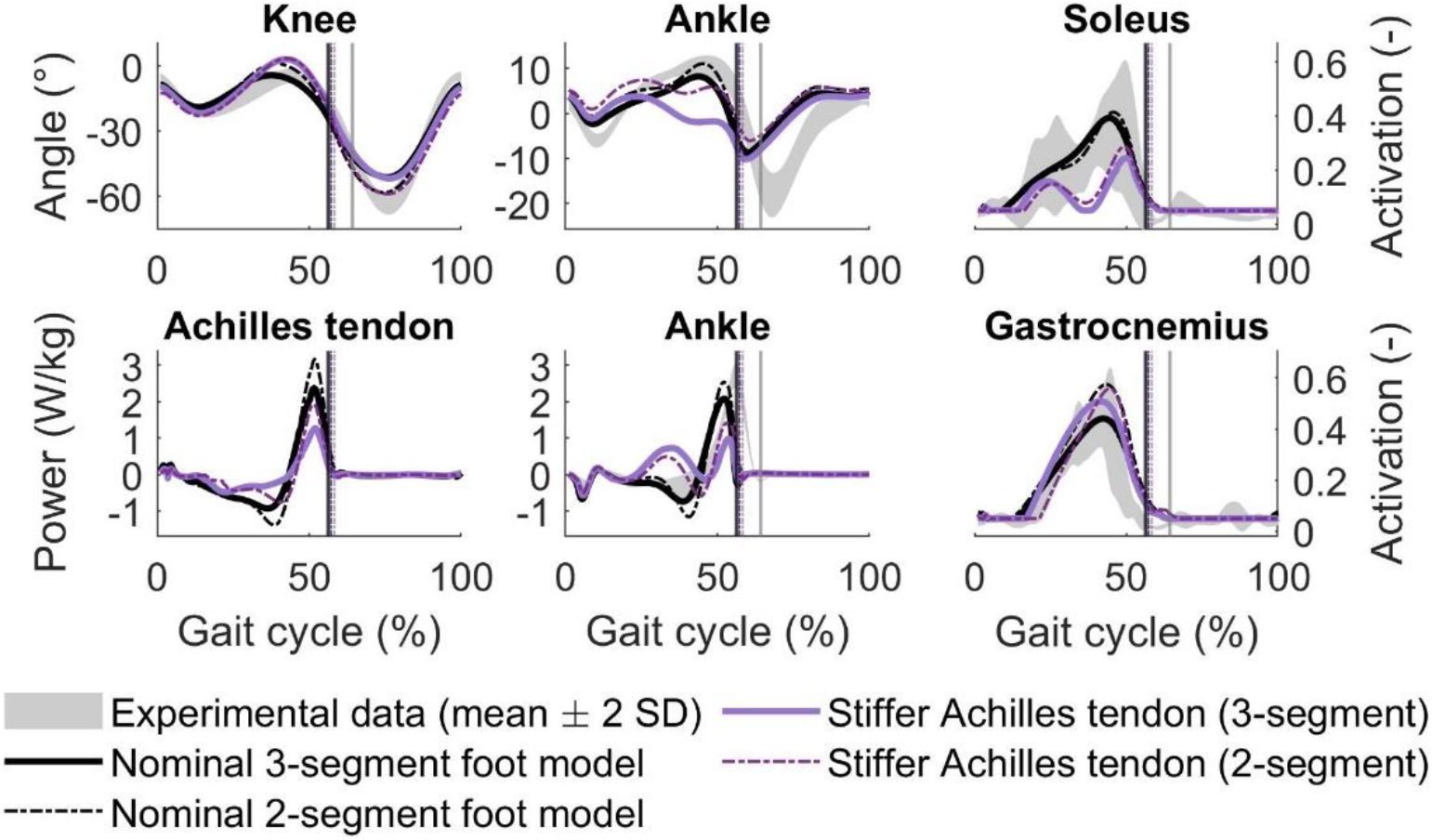
Effect of Achilles tendon stiffness. A stiff Achilles tendon resulted in less ankle dorsiflexion, and more knee extension. Soleus activation decreased, gastrocnemius (medial shown here) activation increased with high tendon stiffness. A stiff Achilles tendon stored less energy and reduced ankle push-off power.

### Plantar fascia and intrinsic foot muscle

Intrinsic foot muscles and a stiff plantar fascia were important to obtain knee extension at push-off and ankle dorsiflexion during mid-stance. Reducing plantar fascia stiffness (i.e. using model Gefen2002 instead of Natali2010) or removing plantar intrinsic muscle results in gait pattern with increased stance knee flexion and increased midtarsal extension (Figure 7). In addition, when the foot is more compliant due to reduced plantar fascia stiffness or absence of the intrinsic foot muscles, we observe that the Achilles tendon stores less elastic energy. This reduced storage and release of elastic energy in the Achilles tendon might explain the increased cost of transport in the models with more compliant feet. These effects were further amplified when using a model with both a compliant plantar fascia and no intrinsic foot muscle. In this model, toe extension was higher than in the nominal 3-segment foot model, and hence the plantar fascia works at higher lengths where it is stiffer due to the nonlinear force-length characteristic. Increased toe extension might be a mechanism to stiffen the foot. In addition, vertical ground reaction force lacks a second peak, showing a reduced push-off in the model with compliant plantar fascia and no intrinsic foot muscles. Remarkably, the simulated gait is not sensitive to parameters of the plantar intrinsic muscle in models with a stiff plantar fascia (supplement S5).

**Figure 7:**
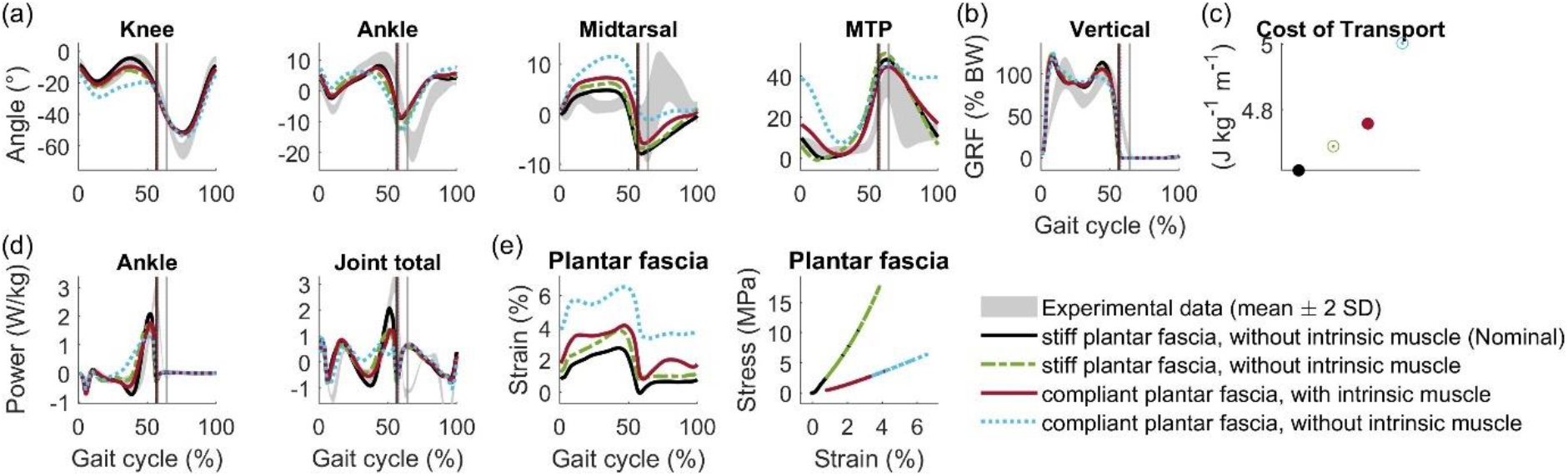
Effect of plantar fascia stiffness and intrinsic muscles. Reducing plantar fascia stiffness and/or removing plantar intrinsic muscle results in gait pattern with flexed stance knee and increased ankle dorsiflexion. Extrinsic toe extensors are more active during stance. They tension the plantar fascia, such that it works on a higher and steeper part of its force-length characteristic. Vertical GRF in late stance, and peak Achilles tendon power decrease, showing a reduced push-off.

### Foot arch height

Simulations based on the 3-segment foot model are very sensitive to foot arch height (Figure 8). Simulations with a smaller arch height, that is not representative for a healthy person, resulted in a gait pattern with increased stance knee flexion and midtarsal extension. The push-off was reduced in the simulations with a smaller arch height in the 3-segment foot model. The reduced push-off is characterised by a reduced vertical ground reaction force at the end of the stance phase and a reduction in total joint power of stance leg. Simulations based on the 2-segment foot model are not sensitive to the foot arch height.

**Figure 8:**
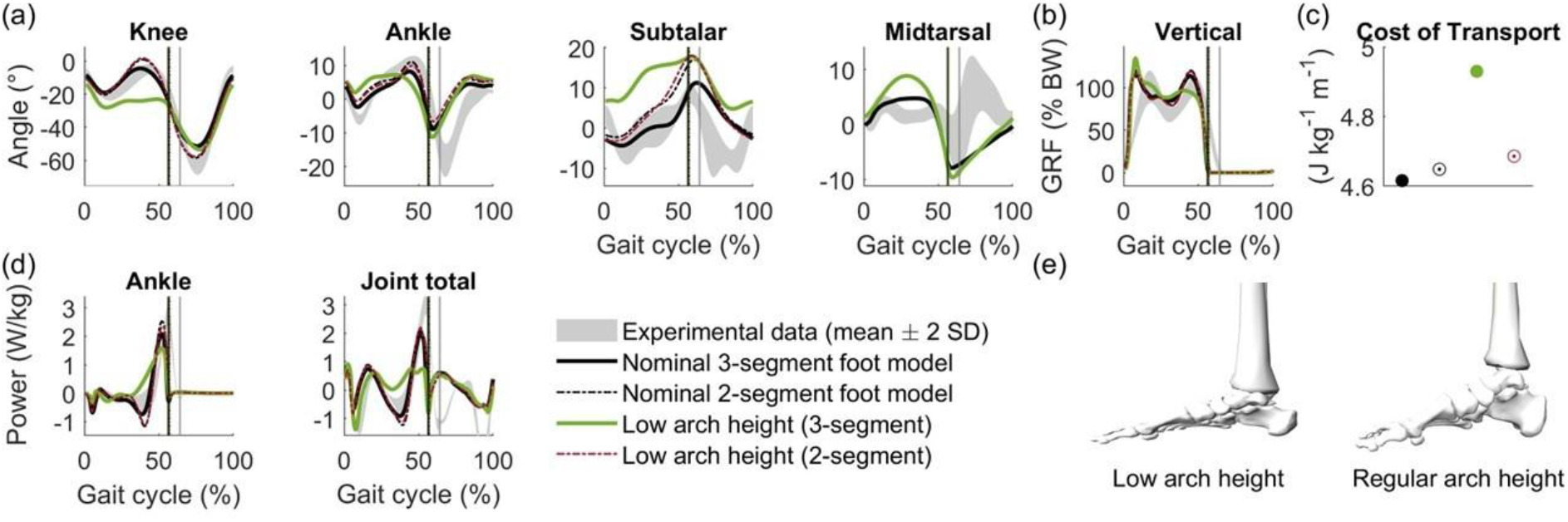
In the 3-segment model, a foot with low arch results in a flexed knee in late stance. The subtalar joint is more inverted, which reduces loading of arch. Midtarsal range of motion during stance is higher, indicating that the arch still deforms a lot more. Vertical GRF lacks 2nd peak, and total joint power of right leg shows no peaks in late stance. There is no real push-off and cost of transport is higher. Arch height had little effect for 2-segment model.

## Discussion

We proposed a novel 3-segment foot model for use in predictive simulations of human locomotion. Simulations based on our model generate physiologically plausible gait mechanics and energetics, and elicited the effect of foot structure on whole body gait mechanics and energetics. (1) The foot-ground contact needs to be sufficiently stiff to obtain mid-stance knee flexion, also known as the loading response. When the foot-ground contact is compliant, negative peak knee joint power is smaller meaning that soft tissues around the knee absorb less of the impact energy from initial contact. (2) The Achilles tendon needs to be sufficiently compliant to store and release strain energy. If the Achilles tendon is stiffer than in humans, the optimal strategy is a gait pattern without ankle dorsiflexion during mid-stance that does not use energy storage and release in the tendon. (3) A stiff plantar fascia, sufficient foot arch height, and intrinsic foot muscles were important to stiffen the foot and generate high ankle push-off powers. With reduced foot stiffness, the optimal gait strategy had reduced ankle push-off power and increased stance knee flexion. (4) During terminal stance plantar fascia and intrinsic foot muscle transfer energy from the MTP to the midtarsal joint to further increase push-off power generated by arch recoil. Our 2-segment model with well-tuned foot-ground contact and Achilles tendon stiffness also captured knee and ankle kinematics, features that had been difficult to predict with previous models, suggesting that a stiff foot is a reasonable approximation for a healthy foot during walking. However, the 2-segment model gives little insights into how different structures contribute to foot stiffness. Furthermore, a more detailed foot model is crucial to investigate how foot pathologies influence whole body mechanics and energetics.

Our simulations confirm that a stiff foot is required to efficiently generate high ankle powers. It has been suggested that the ability to stiffen the foot is important to generate a powerful push-off (24). In simulation, we could easily manipulate foot stiffness by altering arch height, plantar fascia stiffness, or the properties of intrinsic foot muscles. When we reduced the ability to stiffen the foot, simulated push-off ankle power decreased (Figure 7d, 8d), confirming that the ability to stiffen the foot is crucial for efficient push-off. Our simulations suggest that if the foot cannot act as a sufficiently stiff lever, the optimal strategy is to walk with a flexed knee. This is in agreement with the hypothesised contribution of increased foot compliance to crouch gait in children with cerebral palsy (77).

The ability to stiffen the foot is also important for metabolic energy efficiency. Removing the intrinsic foot muscles and reducing plantar fascia stiffness caused a 8.4% increase in the simulated cost of transport in the 3-segment foot model. Similarly, reducing the height of the foot arch (with 1.15 cm) caused a 6.8% increase in the simulated cost of transport in the 3-segment foot model. These alterations in cost of transport cannot be attributed to a few muscles since changes in foot properties affected the metabolic energy expenditure of all muscles (Figure S26). Given that minimising metabolic energy is an important part in our objective function, this suggests that the ability to stiffen the foot is crucial for efficient locomotion. A 2-segment foot, providing a rigid lever, would be expected to be more efficient than a 3-segment foot, but both yield similar costs of transport (Figure 8). While the 2-segment foot model stores more elastic energy in the Achilles tendon than the 3-segment foot model, energy dissipation in the foot itself is four times as high (Figure S25).

Our simulations show that plantar fascia and plantar intrinsic muscle provide push-off power around the midtarsal joint by releasing strain energy and transferring energy absorbed around the MTP joint. Recent experiments indicate that the windlass mechanism contributes to running efficiency by transferring energy from the MTP joint to the arch (78), and by modulating elastic energy storage in the arch (79). Measuring individual contributions to energy storage and transfer by plantar fascia, intrinsic muscles, and extrinsic muscles is impossible experimentally. However in simulations, this information is readily available. In the 3-segment foot model, the plantar fascia and intrinsic muscle absorb most braking energy at the MTP joint and transfer this to the midtarsal joint (Figure 9). Releasing strain energy from the plantar fascia and the tendon of the intrinsic muscle, and active contraction of the intrinsic muscle provide additional power to the midtarsal joint (Figure 9). The 2-segment foot model also absorbs energy around the MTP (Figure 9) but the energy is dissipated (supplement S16), thus a rigid foot arch does not yield the most efficient push-off. Interestingly, the energy efficient mechanisms that we find in walking simulations with a 3-segment foot model, are the same mechanisms as experimentally observed in running (78,79).

**Figure 9:**
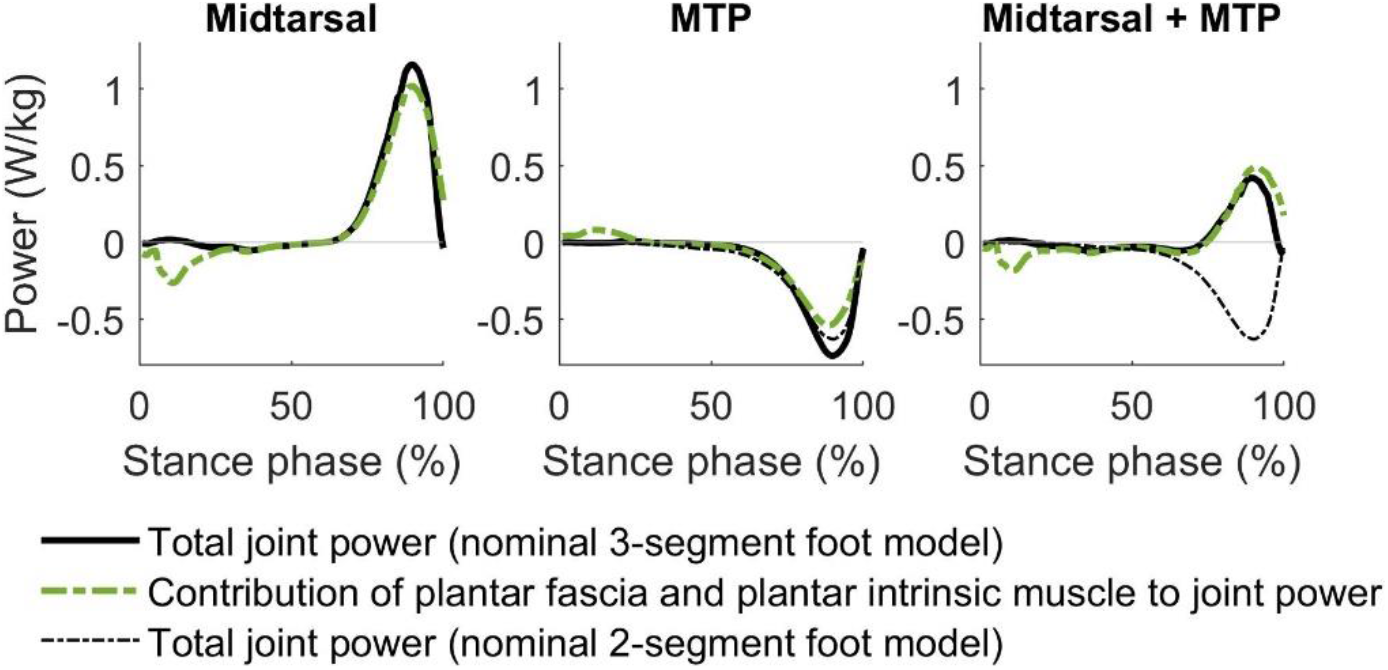
Joint power of midtarsal joint, MTP joint, and summed power of both joints. Plantar fascia and plantar intrinsic muscle account for most power around these joints. Separate contributions of plantar fascia and intrinsic muscle have similar magnitude (Figure S30).

Our simulations support the idea that efficiency of upright walking might have driven the evolution towards a stiffer foot. The simulated gait based on a model with a low arch and without plantar fascia has striking resemblances to bipedal chimpanzee gait. Due to the absence of a foot arch and a small plantar fascia, chimpanzees are thought to be unable to stiffen their foot through the windlass mechanism (22,80). They are able to stiffen their foot by muscle contraction, but cannot reach human levels of foot stiffness (22). We modelled this conceptually by a low-arched foot without plantar fascia. The corresponding gait pattern has higher knee flexion and ankle dorsiflexion than our nominal model, similar – yet less pronounced - to how bipedal chimpanzee and human walking differ (Figure 10a). Consistent with experimental observations in chimpanzees, this extreme reduction in foot stiffness resulted in a knee extension moment throughout the stance phase, a reduced peak ankle plantarflexion moment, reduced forward ground reaction forces during terminal stance, and the absence of a second peak in the vertical ground reaction force (Figure 10b,c). It is interesting how only altering foot properties induces many features of bipedal chimpanzee locomotion given that there are many more morphological differences between humans and chimpanzees, such as for example the longer plantar flexor muscle fibres and shorter Achilles tendon in chimpanzees than in humans.

**Figure 10:**
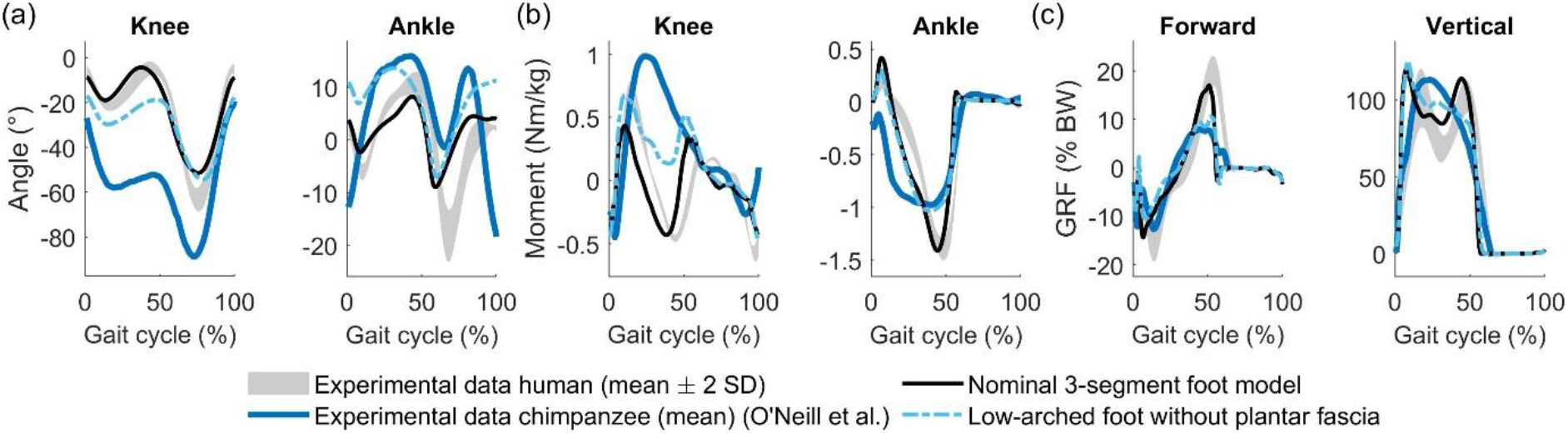
Comparing simulated gait to experimental results of bipedal chimpanzee walking. Chimpanzee data was digitised from O’Neill et al. (81,82). (a) Kinematics. (b) Kinetics. We scaled dimensionless moments by multiplying with leg length of our reference subject (85 cm). (c) Ground reaction forces (GRF) expressed as percentage of body weight (%BW).

Our simulations only partially capture the effect of inhibiting intrinsic foot muscle activation on whole-body gait mechanics and energetics, indicating that further modelling refinements might be needed. Farris et al. applied a nerve block to the plantar intrinsic foot muscles (31). We mimicked this experiment in simulation by imposing a low constant activation to the plantar intrinsic muscles (for detailed results, see supplement S14). Consistent with experiments (31), limiting intrinsic muscle activity in simulation reduced ankle-foot push-off power, increased hip work (Figure S29), and hardly affected cost of transport (+1.2%). In contrast to experimental data, limiting intrinsic muscle activity in simulation increased arch compression (Figure S27). This suggest that our model overestimates the contribution of active muscle force to arch stiffness. The moment arm of our lumped plantar intrinsic muscle around the midtarsal joint was based on the moment arm of the flexor digitorum brevis, whereas the abductor hallucis and quadratus plantae muscles, which are not represented separately in our model, have smaller moment arms (26). However, when we reduced the ability of the intrinsic foot muscle to stiffen the foot arch, our simulations of healthy walking became less realistic. The high contribution of active force of the intrinsic foot models might compensate for stiffening mechanisms that we did not model. Due to the shape of the tarsal bones (19), we expect that even small levels of muscle co-contraction will cause them to interlock, thereby increasing stiffness. Mann and Inman did indeed observe co-contraction from plantar intrinsic muscles and extensor digitorum brevis in late stance (30).

## Data accessibility

Code and models: https://github.com/Lars-DHondt-KUL/3dpredictsim

Simulation results: will be made available

## Acknowledgements

We thank Madhu Venkadesan for the useful discussions about foot mechanics and energetics.

