## Supplementary material for "A dynamic foot model for predictive simulations of gait reveals causal relations between foot structure and whole body mechanics"

### S1. Achilles tendon stiffness

Table S1 Optimal Achilles tendon stiffness.
Preliminary simulations indicated that reducing the normalised stiffness of the Achilles tendon to 50% of its value results in the lowest optimal cost. Evaluating our choice with the final model (and finer mesh density) shows that 50% and 60% stiffness have comparable cost, thus confirming our choice.

| Achilles tendon stiffness [%] | Optimal cost (preliminary) | Optimal cost (final) |
|---|---|---|
| 30 | 372.4996 | 359.4623 |
| 40 | 359.1847 | 344.0870 |
| <u>50</u> | <u>356.6144</u> | 341.6931 |
| 60 | 358.7890 | 341.5860 |
| 70 | 361.9741 | 345.0313 |
| 80 | 360.4416 | 345.8073 |
| 90 | 363.5739 | 344.8347 |
| 100 | 362.1360 | 347.8070 |

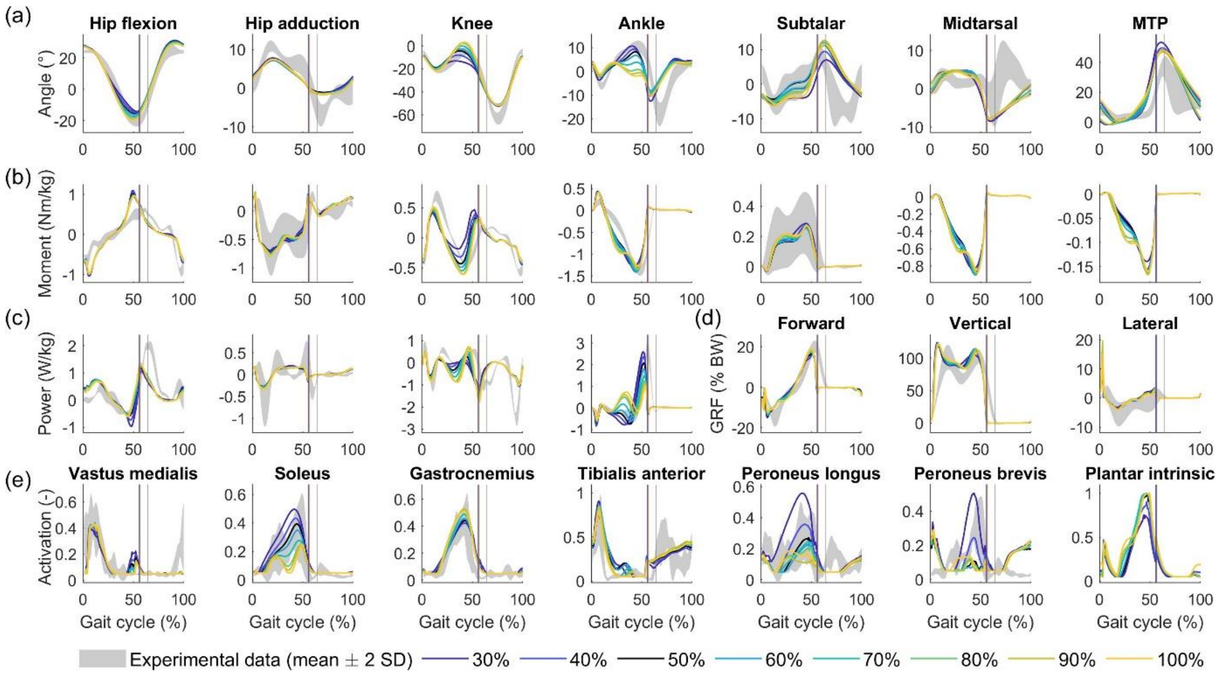


Figure S1 Effect of Achilles tendon stiffness. Normalised stiffness is reduced to a percentage of the reference value (1). (a) Kinematics. (b) Kinetics. (c) Joint powers. (d) Ground reaction forces, expressed as % body weight. (e) Muscle activation. Gastrocnemius indicates the medial gastrocnemius.

### S2. Passive isometric ankle moment

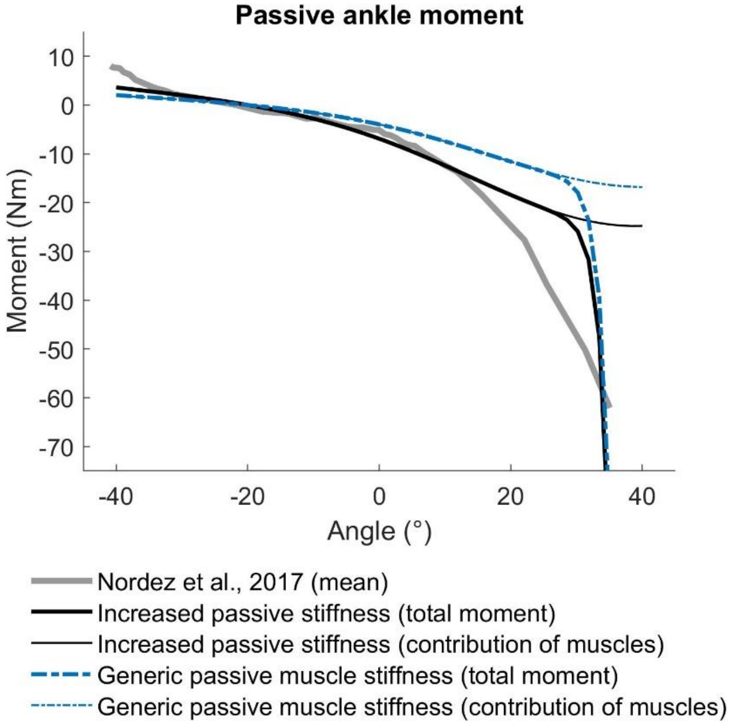


Figure S2 Passive isometric ankle moments. Dorsiflexion is positive. Thin lines show the contribution of muscles. Thick lines show total, also including coordinate limit torques (2). Reference data was digitised from Nordez et al (3).

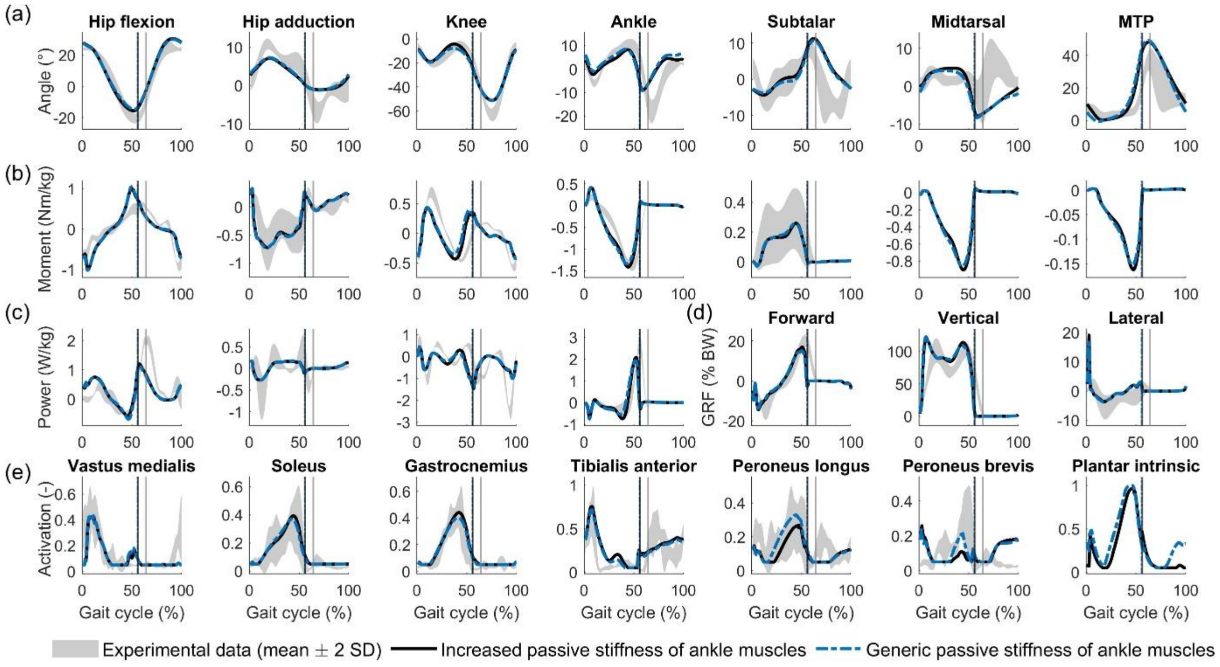


Figure S3 Effect of shifting the passive force-length characteristic of all muscles crossing the ankle towards 10% shorter normalised fibre lengths (1). (a) Kinematics. (b) Kinetics. (c) Joint powers. (d) Ground reaction forces, expressed as % body weight. (e) Muscle activation. Gastrocnemius indicates the medial gastrocnemius.

### S3. Triceps surae maximal isometric force

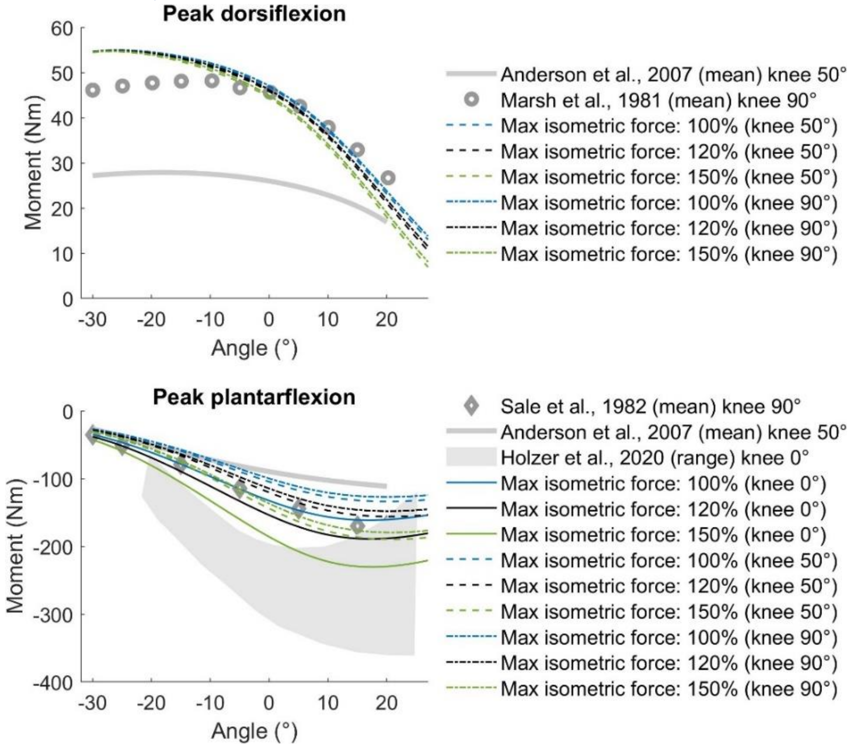


Figure S4 Effect of triceps surae maximal isometric force on maximal isometric ankle moments. Dorsiflexion is positive. Knee flexion angles were adjusted to match experimental conditions. Experimental data taken from (4–7).

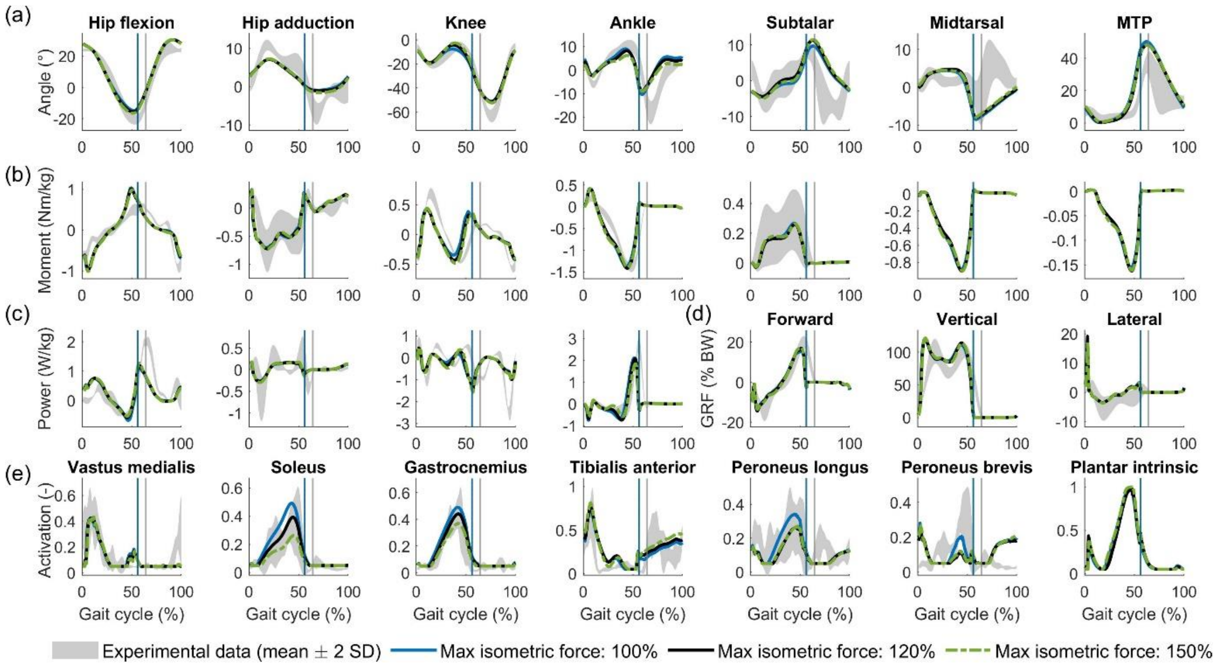


Figure S5 Effect of triceps surae maximal isometric force. (a) Kinematics. (b) Kinetics. (c) Joint powers. (d) Ground reaction forces, expressed as % body weight. (e) Muscle activation. Gastrocnemius indicates the medial gastrocnemius.

#### S4. Midtarsal joint axis orientation

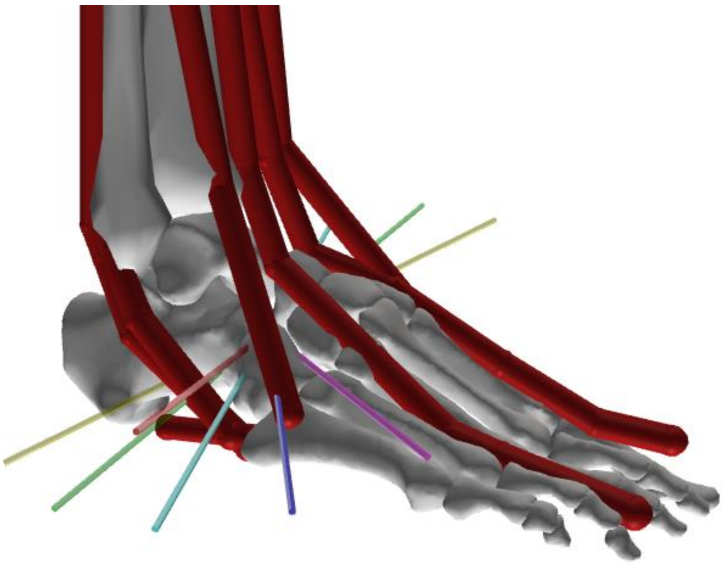

Figure S6 Orientation of the different midtarsal joint axes considered. Colours are consistent with Figure S7 and Figure S8.

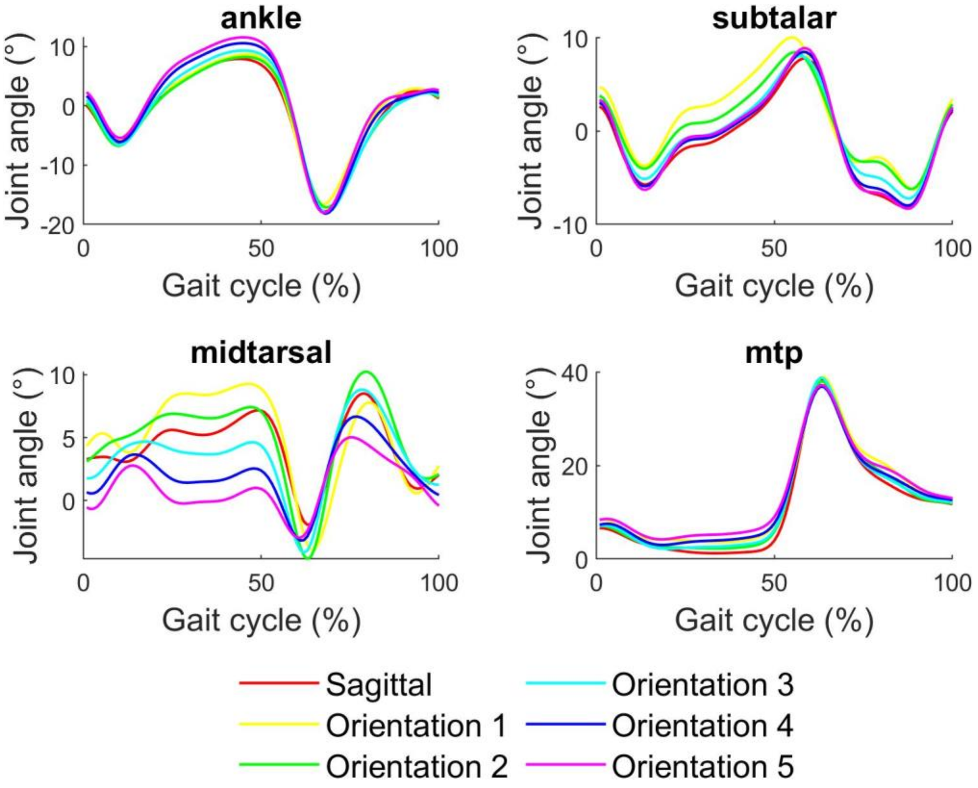


Figure S7 Effect of midtarsal joint axis orientation on inverse kinematics. Mean inverse kinematics of 10 strides overground walking at self-selected speed (1.33 m $s^{-1}$). Proximal joints are not shown because there was no considerable effect.

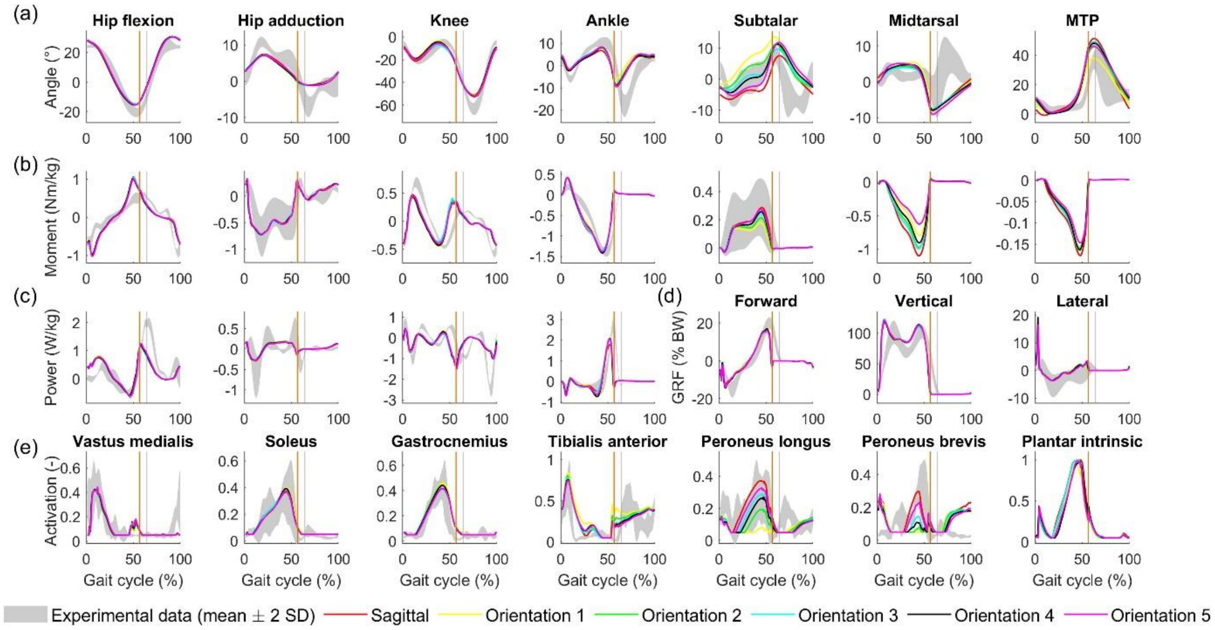


Figure S8 Effect of midtarsal joint axis orientation on simulated gait. (a) Kinematics. (b) Kinetics. (c) Joint powers. (d) Ground reaction forces, expressed as % body weight. (e) Muscle activation. Gastrocnemius indicates the medial gastrocnemius.

### S5. Plantar intrinsic muscle parameter sensitivity

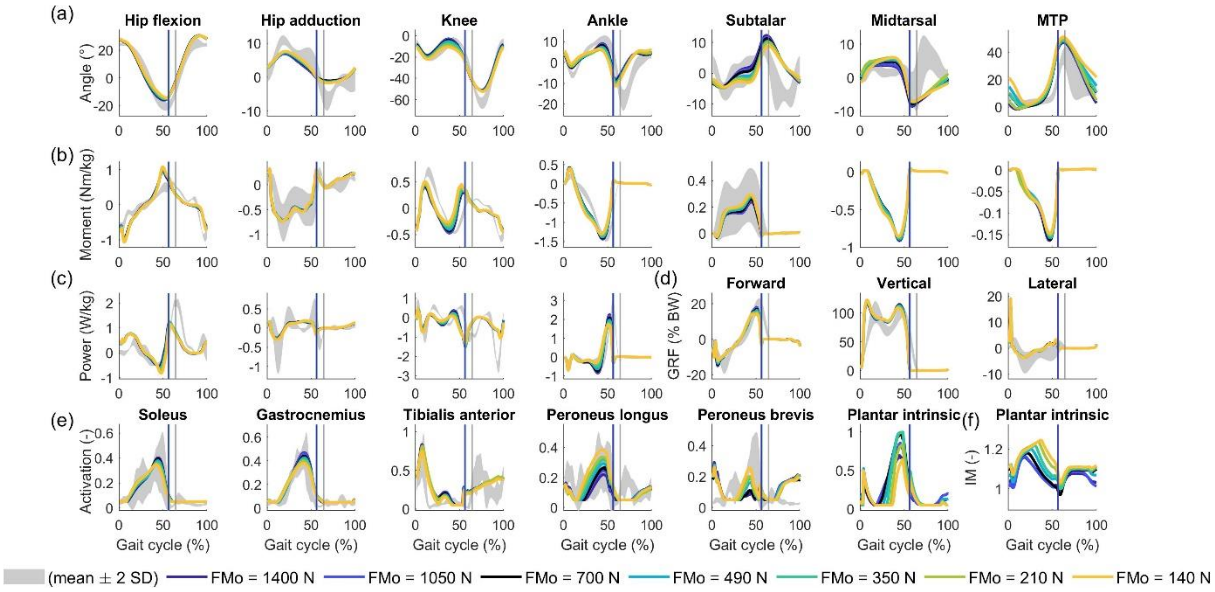


Figure S9 Effect of plantar intrinsic muscle maximal isometric force (FMo) on simulated gait.
(a) Kinematics. (b) Kinetics. (c) Joint powers. (d) Ground reaction forces, expressed as % body weight. (e) Muscle activation. Gastrocnemius indicates the medial gastrocnemius. (f) Fibre length, normalised to optimal fibre length.

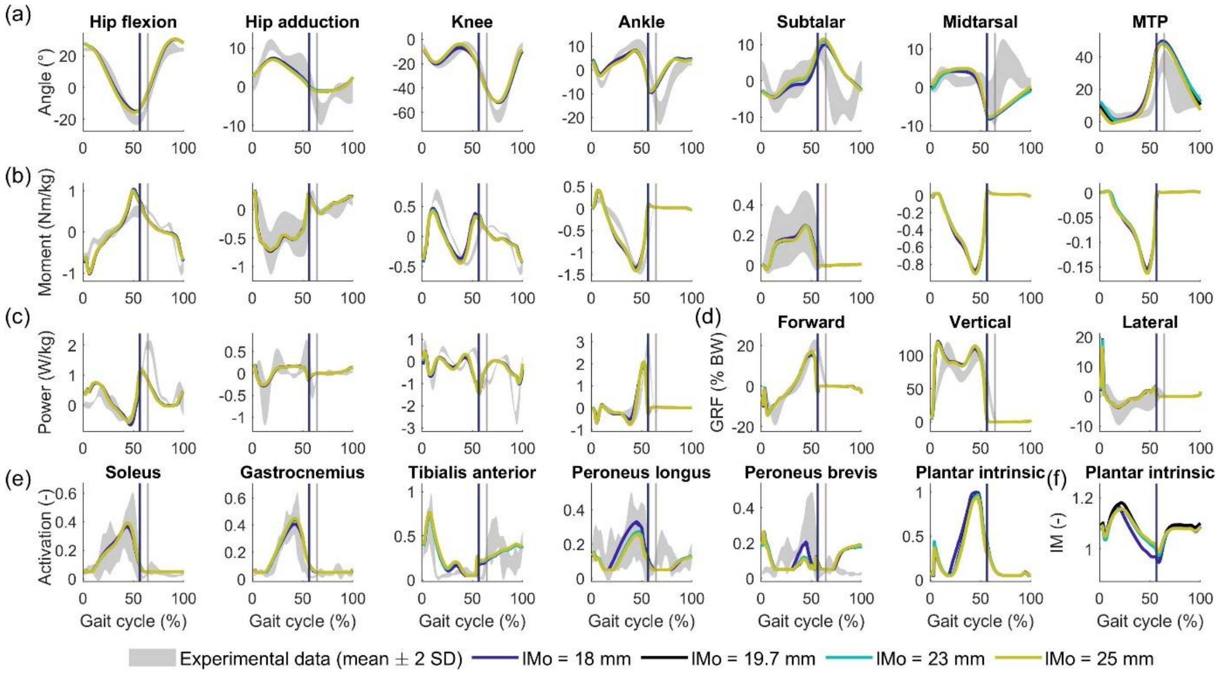


Figure S10 Effect of plantar intrinsic muscle optimal fibre length (lMo) on simulated gait.
(a) Kinematics. (b) Kinetics. (c) Joint powers. (d) Ground reaction forces, expressed as % body weight. (e) Muscle activation. Gastrocnemius indicates the medial gastrocnemius. (f) Fibre length, normalised to optimal fibre length.

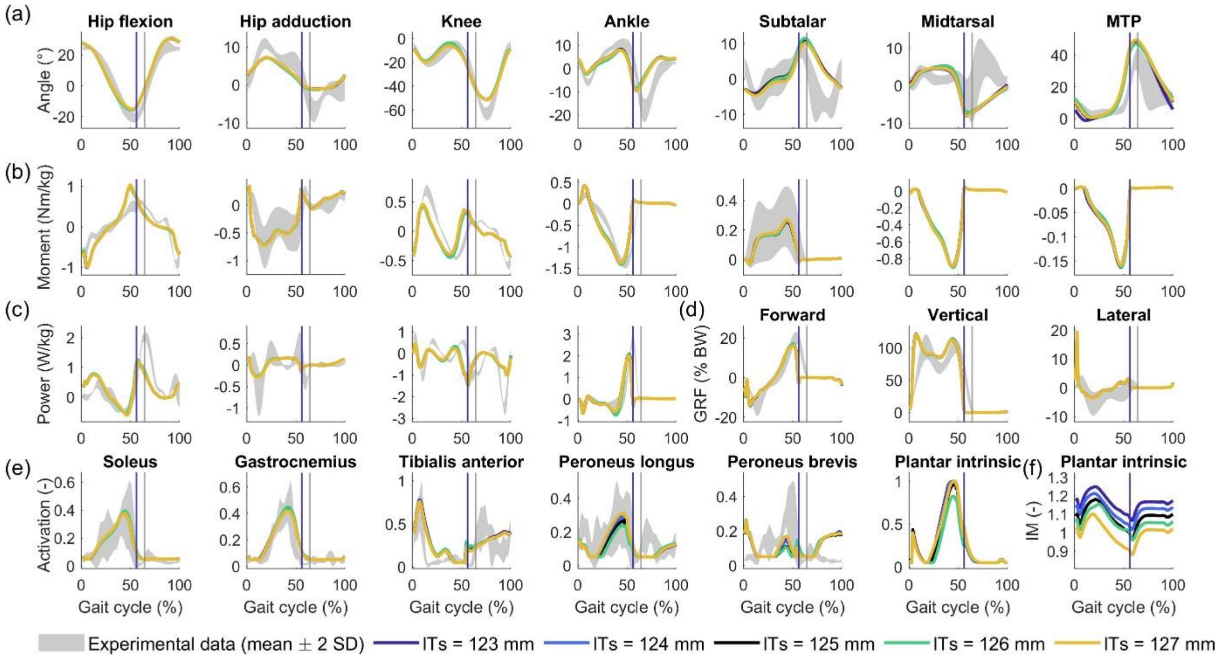


Figure S11 Effect of plantar intrinsic muscle maximal isometric force (FMo) on simulated gait. (a) Kinematics. (b) Kinetics. (c) Joint powers. (d) Ground reaction forces, expressed as % body weight. (e) Muscle activation. Gastrocnemius indicates the medial gastrocnemius. (f) Fibre length, normalised to optimal fibre length.

### S6. Approximating conditional statements with a hyperbolic tangent

Elaborate explanation of smoothing is provided in the supplementary material of (8).

The metabolic energy model (9) contains terms that are not continuously differentiable, such as the work rate of a muscle fibre

$$\dot{W} = \begin{cases} -F^T * v^M \text{ , } if\ v^M \leq 0 \\ 0 \text{ , } if\ v^M > 0 \end{cases}$$

To make this model compatible with algorithmic differentiation, it was approximated as

$$\dot{W} = -F^T * v^M * (0{,}5 \ - 0{,}5 \tanh(b * v^M))$$

$$\dot{W} = -F^T * v^M_{negative}$$

The smoothing coefficient (b) determines how smooth the transition between the conditions is. A lower value results in a smoother curve, however also lead to distortion in a larger range of values around 0 (see figure). Since using b = 10 (cfr. (2)) leads to distortion of the negative contraction velocities, and thus an underrepresentation of work rate in metabolic energy rate, we opted to use b = 100.

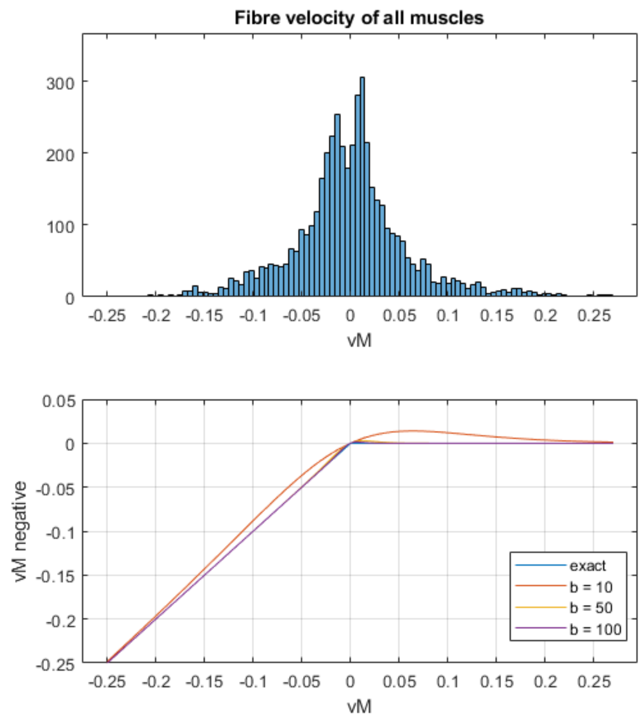


Figure S12 A low smoothing coefficient (b = 10) distorts the negative contraction velocities that are used to calculate work rate of the muscle fibre.

Figure S13 Effect of smoothing coefficient on gait.

Figure S14 Effect of smoothing coefficient on metabolic energy.

#### S7. Convergence analysis

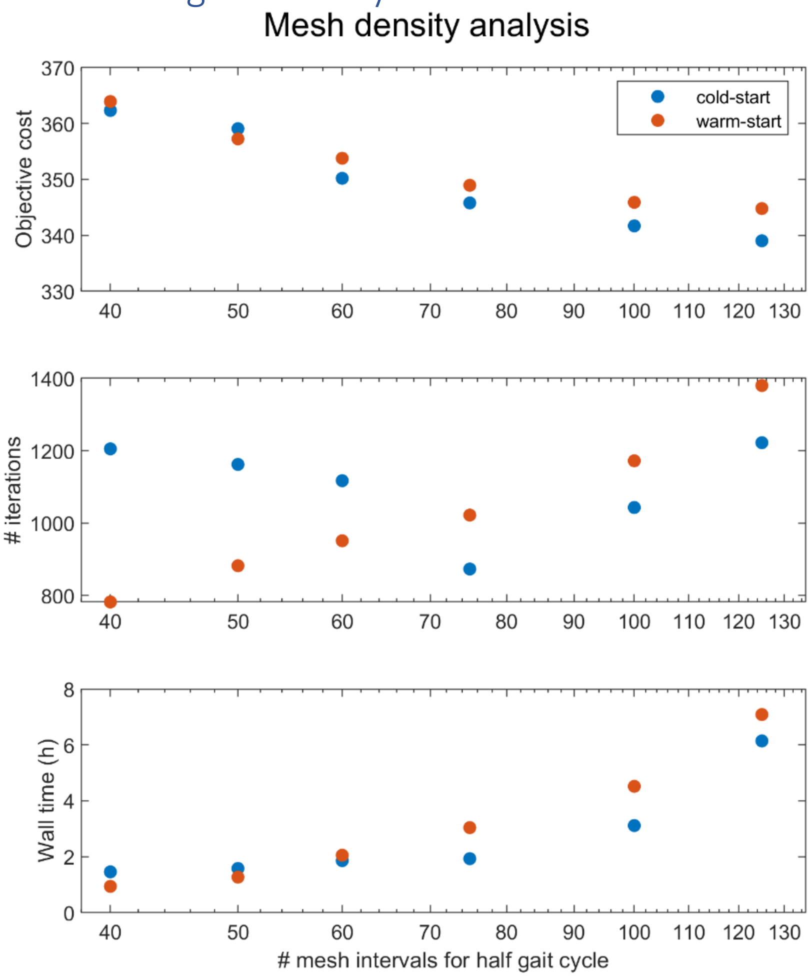


Figure S15 We simulated gait for the nominal 3-segment foot model with different amounts of time mesh intervals and initial guesses. Cold-start (i.e. without any reference gait data) resulted in lower objective cost overall. Refining the mesh further than 100 intervals did not yield a relevant improvement in objective cost (1% improvement), thus we selected 100 mesh intervals.

*Table S2 Convergence analysis.*
*Reducing convergence tolerance of the optimisation solver below 1E-04 has a negligible effect on optimal cost.*

| IPOPT tolerance | Optimal cost |
|---|---|
| 1E-04 | 341.693 |
| 1E-05 | 341.506 |
| 1E-06 | 341.506 |

#### S8. Marker protocol

*Table S3 Marker placement for the feet.*
*The marker protocol is a subset of the markers used by Boey et al. (10).*

| Segment | Marker | Anatomical position |
|---|---|---|
| Hindfoot | Hindfoot 1 | Craniodorsal calcaneus |
| | Hindfoot 2 | Caudal dorsal calcaneus |
| | Hindfoot 3 | Ventrolateral calcaneus |
| | Hindfoot 4 | Ventromedial calcaneus |
| Forefoot | Forefoot 1 | Basis of fifth metatarsal |
| | Forefoot 3 | Basis of first metatarsal |
| | Forefoot 4 | Head of fifth metatarsal |
| | Forefoot 6 | Head of first metatarsal |
| Toes | Toes 1 | Hallux |
| | Toes 2 | Third toe |

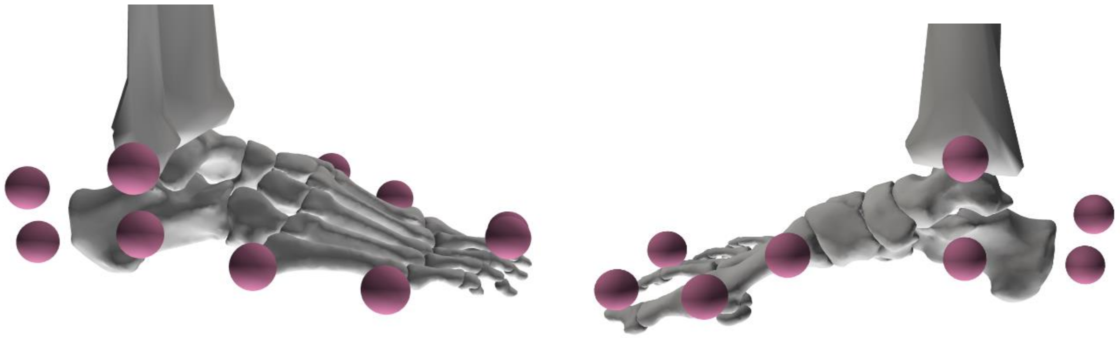

Figure S16 Visualisation of all foot markers (Table S3) and markers on the malleoli.

### S9. Predictive gait simulations with 3-segment and 2-segment foot models

*Table S4 Correlation coefficients joint kinematics*

| | knee | | ankle | | subtalar | | midtarsal | | MTP | |
|---|---|---|---|---|---|---|---|---|---|---|
| | Stance | Swing | Stance | Swing | Stance | Swing | Stance | Swing | Stance | Swing |
| 2-segment foot model Falisse et al. (8) | 0.90 | 0.96 | 0.11 | 0.98 | 0.93 | -0.12 | n/a | n/a | 0.81 | 0.98 |
| 2-segment foot model | 0.87 | 0.91 | 0.88 | 0.99 | 0.87 | 0.18 | n/a | n/a | 0.91 | 0.97 |
| 3-segment foot model | 0.93 | 0.87 | 0.88 | 0.99 | 0.87 | 0.02 | 0.80 | -0.38 | 0.99 | 0.82 |

*Table S5 Correlation coefficients joint kinetics*

| | knee | | ankle | | subtalar | |
|---|---|---|---|---|---|---|
| | Stance | Swing | Stance | Swing | Stance | Swing |
| 2-segment foot model Falisse et al. (8) | 0.95 | 0.91 | 0.95 | 0.40 | 0.63 | -0.19 |
| 2-segment foot model | 0.93 | 0.89 | 0.92 | 0.79 | 0.82 | -0.04 |
| 3-segment foot model | 0.92 | 0.85 | 0.93 | 0.79 | 0.70 | 0.06 |

*Table S6 Correlation coefficients joint powers*

| | knee | | ankle | | subtalar | |
|---|---|---|---|---|---|---|
| | Stance | Swing | Stance | Swing | Stance | Swing |
| 2-segment foot model Falisse et al. (8) | 0.81 | 0.73 | 0.63 | 0.34 | 0.62 | 0.37 |
| 2-segment foot model | 0.86 | 0.66 | 0.72 | 0.80 | 0.71 | 0.18 |
| 3-segment foot model | 0.93 | 0.64 | 0.75 | 0.83 | 0.10 | -0.46 |

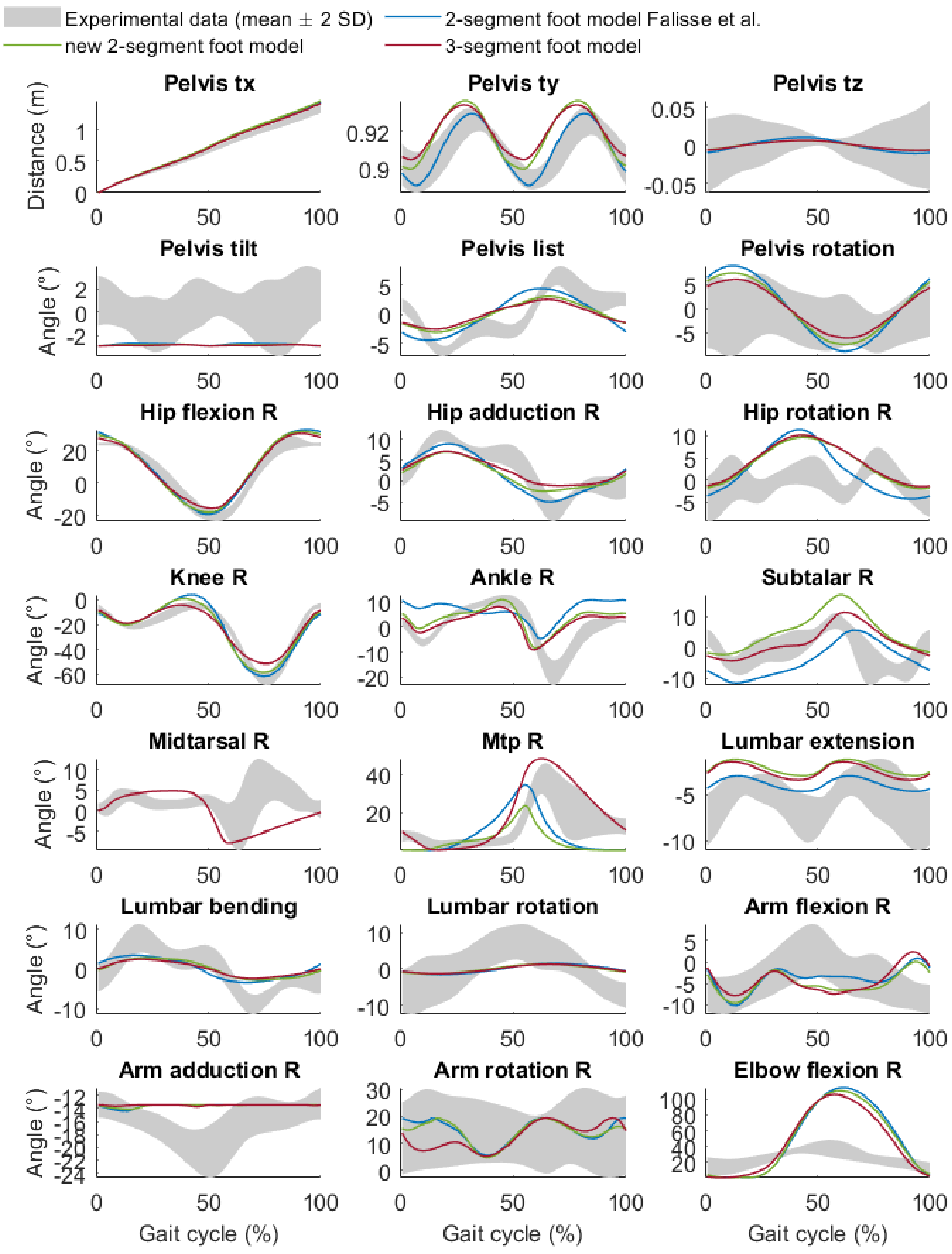


Figure S17 All kinematics (right side) for gait simulations with our nominal 3-segment foot model, 2-segment foot model, and the model from Falisse et al. (8)

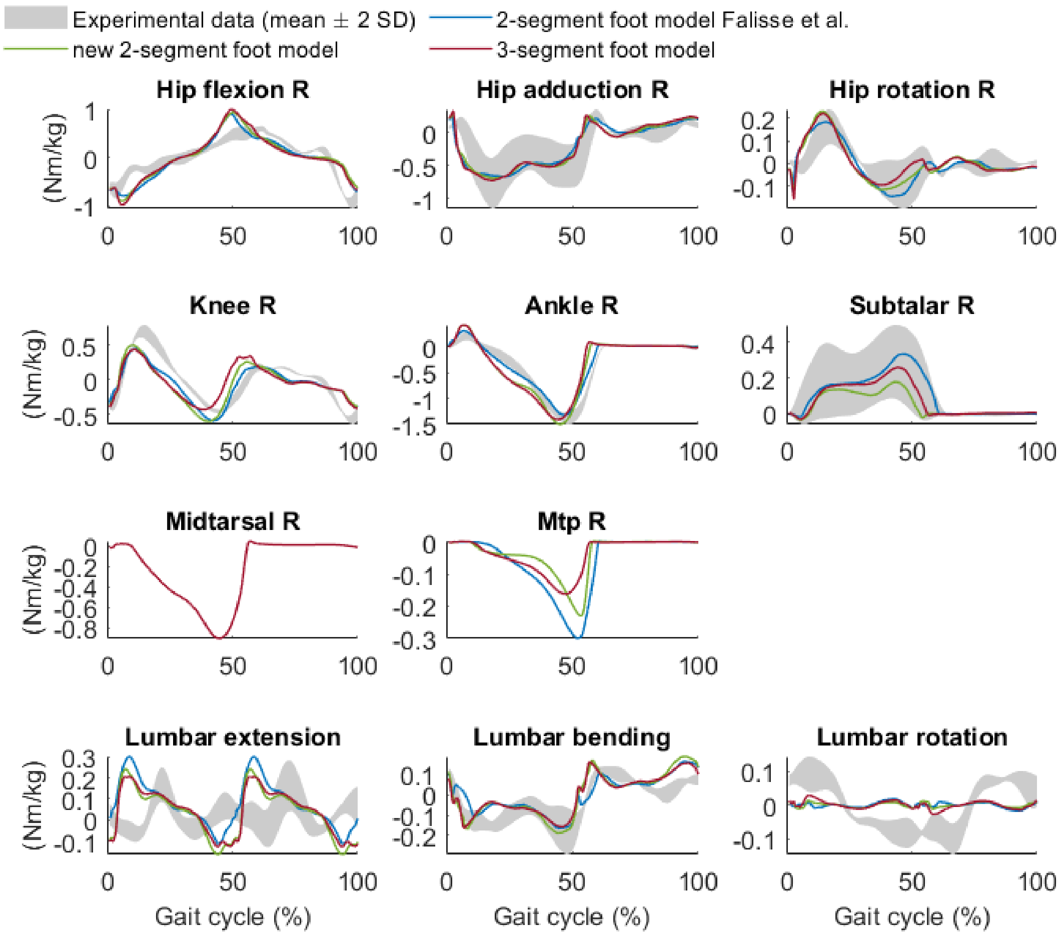


Figure S18 Joint moments of muscle-driven joints (right side) for gait simulations with our nominal 3-segment foot model, 2-segment foot model, and the model from Falisse et al. (8)

#### S10. Unified deformable model as validation

We selected powers calculated with a unified deformable model as reference data, because this method does not depend on rigid-body assumptions or predefined joints (11), thus better captures the true magnitude of power (11). Deformation power distal to a reference segment is calculated based on absolute kinematics of the reference segment and distal ground reaction forces (11). We calculated simulated power as the sum of joint powers and contact deformation powers that are distal to each corresponding segment in our model. Distal to hallux only consists of the contact sphere under the toes. Distal to forefoot includes distal to hallux and also MTP joint and contact spheres under forefoot. Distal to hindfoot includes MTP and midtarsal joints and all contact spheres. Distal to shank is the sum of distal to hindfoot and ankle and subtalar joint powers. Work was calculated by integrating power over the stance phase.

### S11. Plantar fascia stiffness

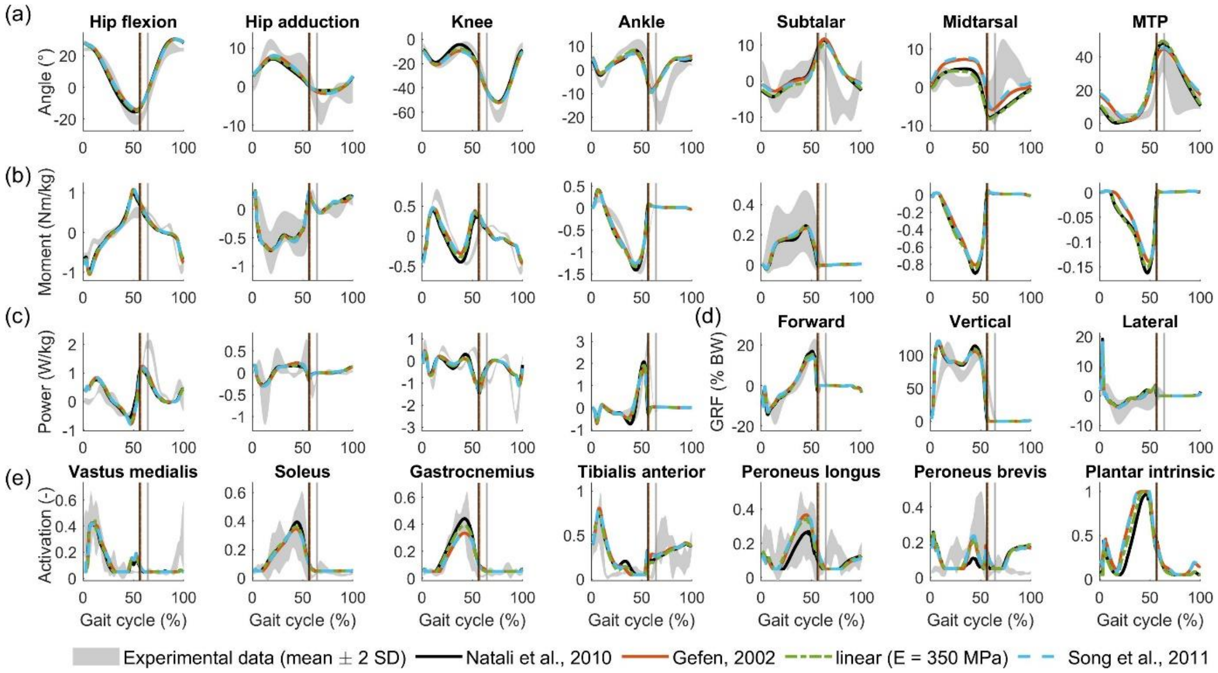


Figure S19 Effect of plantar fascia stiffness on simulated gait. Plantar fascia stiffness models taken from Natali et al. (12), Gefen (13),Young's modulus 350 MPa (14), and Song et al. (15). (a) Kinematics. (b) Kinetics. (c) Joint powers. (d) Ground reaction forces, expressed as % body weight. (e) Muscle activation. Gastrocnemius indicates the medial gastrocnemius.

### S12. Foot-ground contact

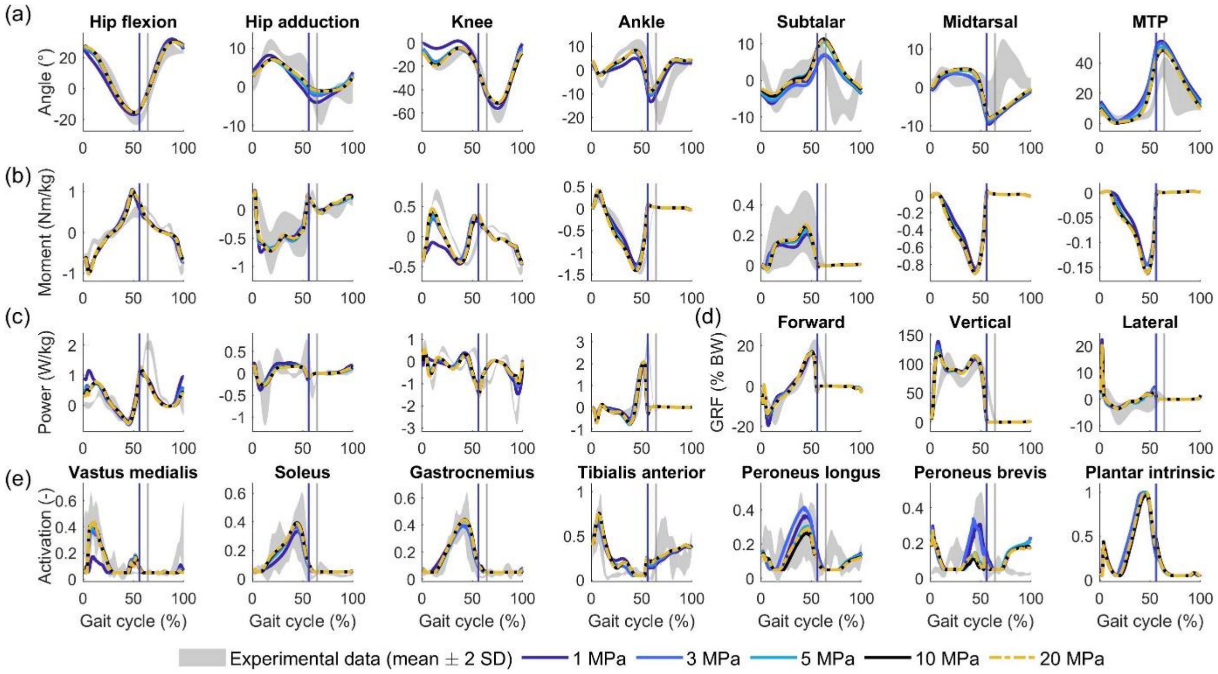


Figure S20 Effect of foot-ground contact stiffness (16) on simulated gait. (a) Kinematics. (b) Kinetics. (c) Joint powers. (d) Ground reaction forces, expressed as % body weight. (e) Muscle activation. Gastrocnemius indicates the medial gastrocnemius.

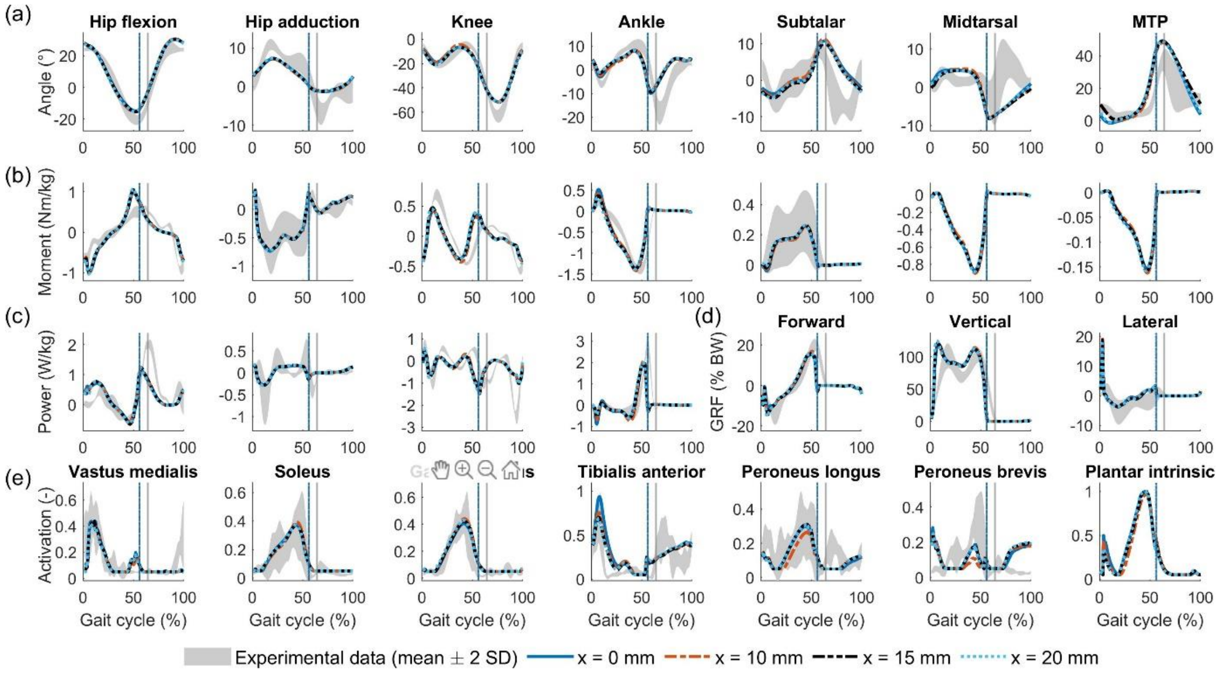


Figure S21 Effect of heel contact sphere position on simulated gait with 3-segment foot model. The nominal models consider the sphere centre at x = 15 mm. (a) Kinematics. (b) Kinetics. (c) Joint powers. (d) Ground reaction forces, expressed as % body weight. (e) Muscle activation. Gastrocnemius indicates the medial gastrocnemius.

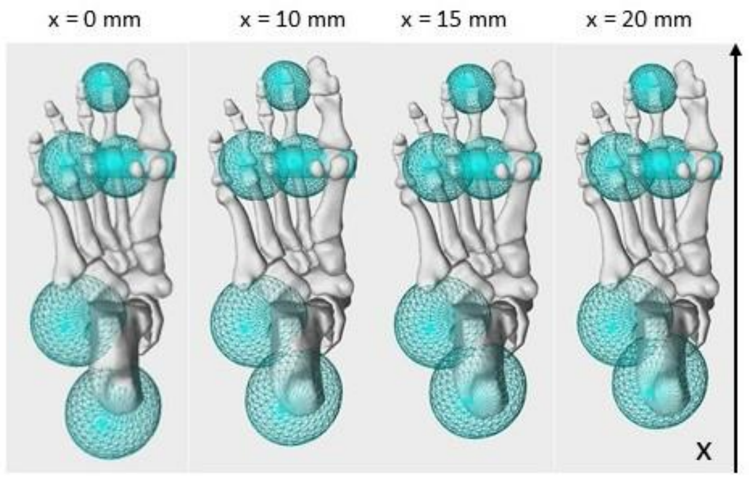


Figure S22 X-position of the heel contact sphere used for simulations in Figure S21. The nominal models consider the sphere centre at x = 15 mm.

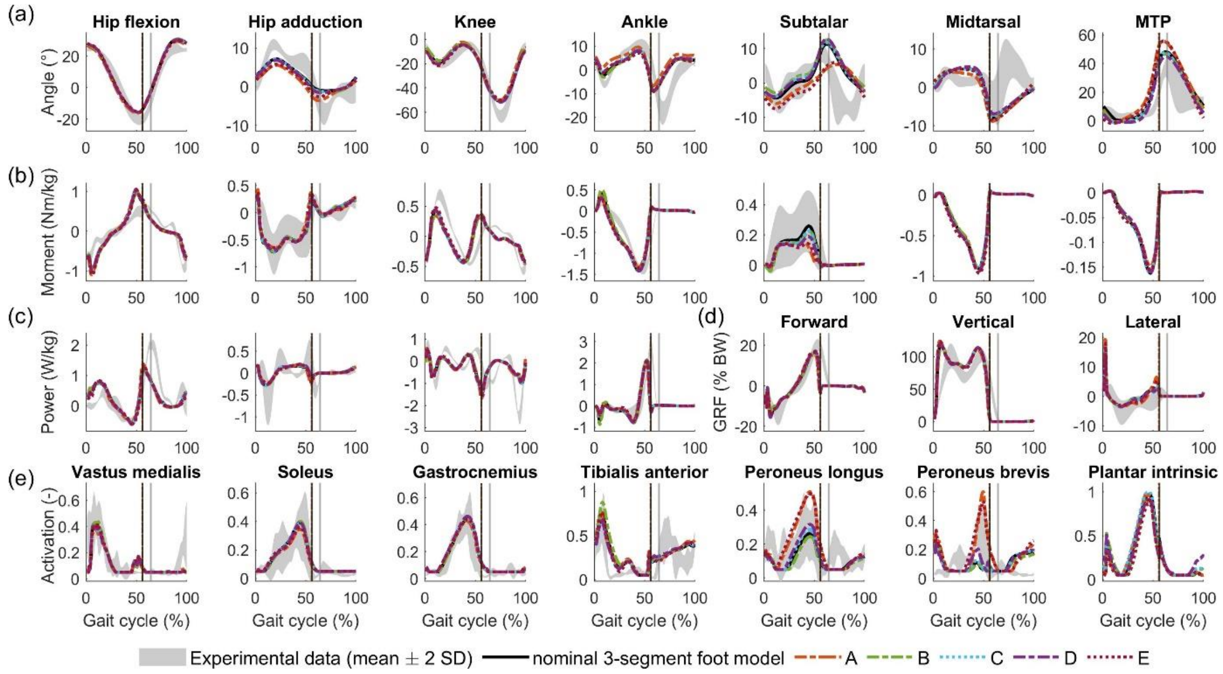


Figure S23 Effect of contact sphere configuration on simulated gait with 3-segment foot model. The nominal models consider the sphere centre at x = 15 mm. (a) Kinematics. (b) Kinetics. (c) Joint powers. (d) Ground reaction forces, expressed as % body weight. (e) Muscle activation. Gastrocnemius indicates the medial gastrocnemius.

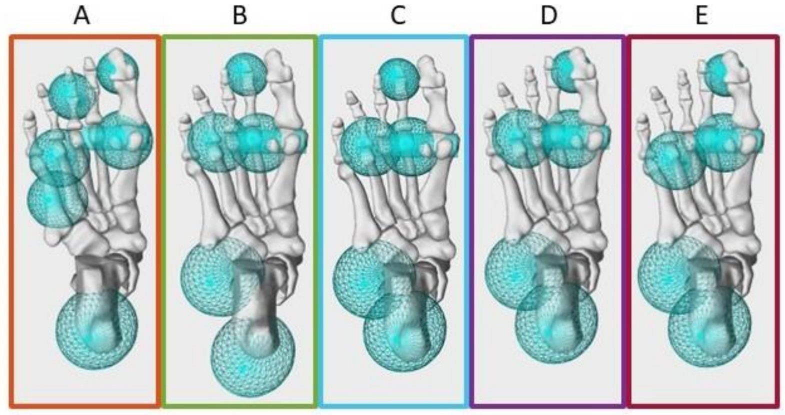


Figure S24 Contact sphere configurations used for simulations in Figure S23.

#### S13. Foot model affects whole-body energetics

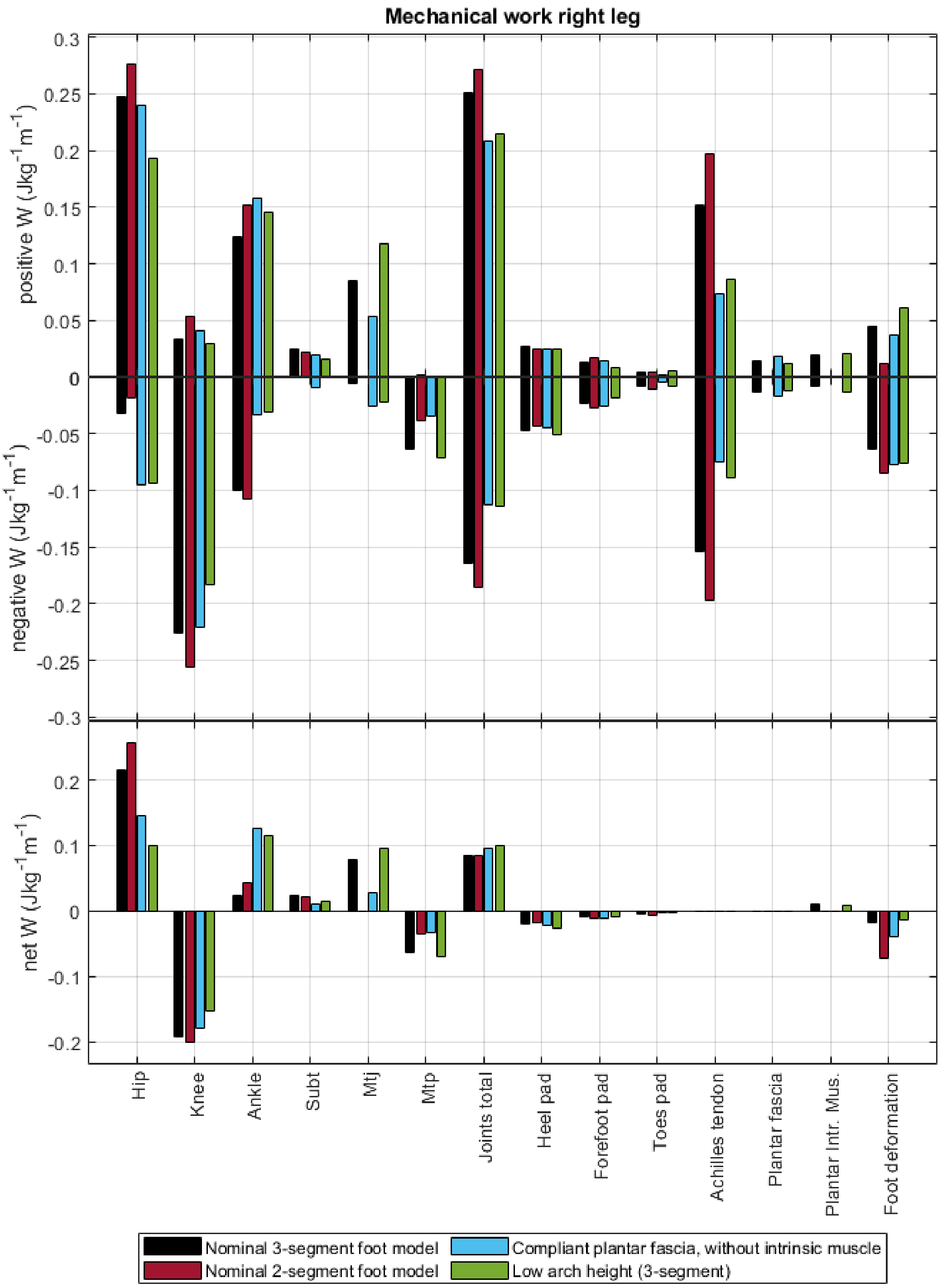


Figure S25 Positive, negative, and net work around joints and by selected structures. Positive net work indicates energy generation, negative energy dissipation.

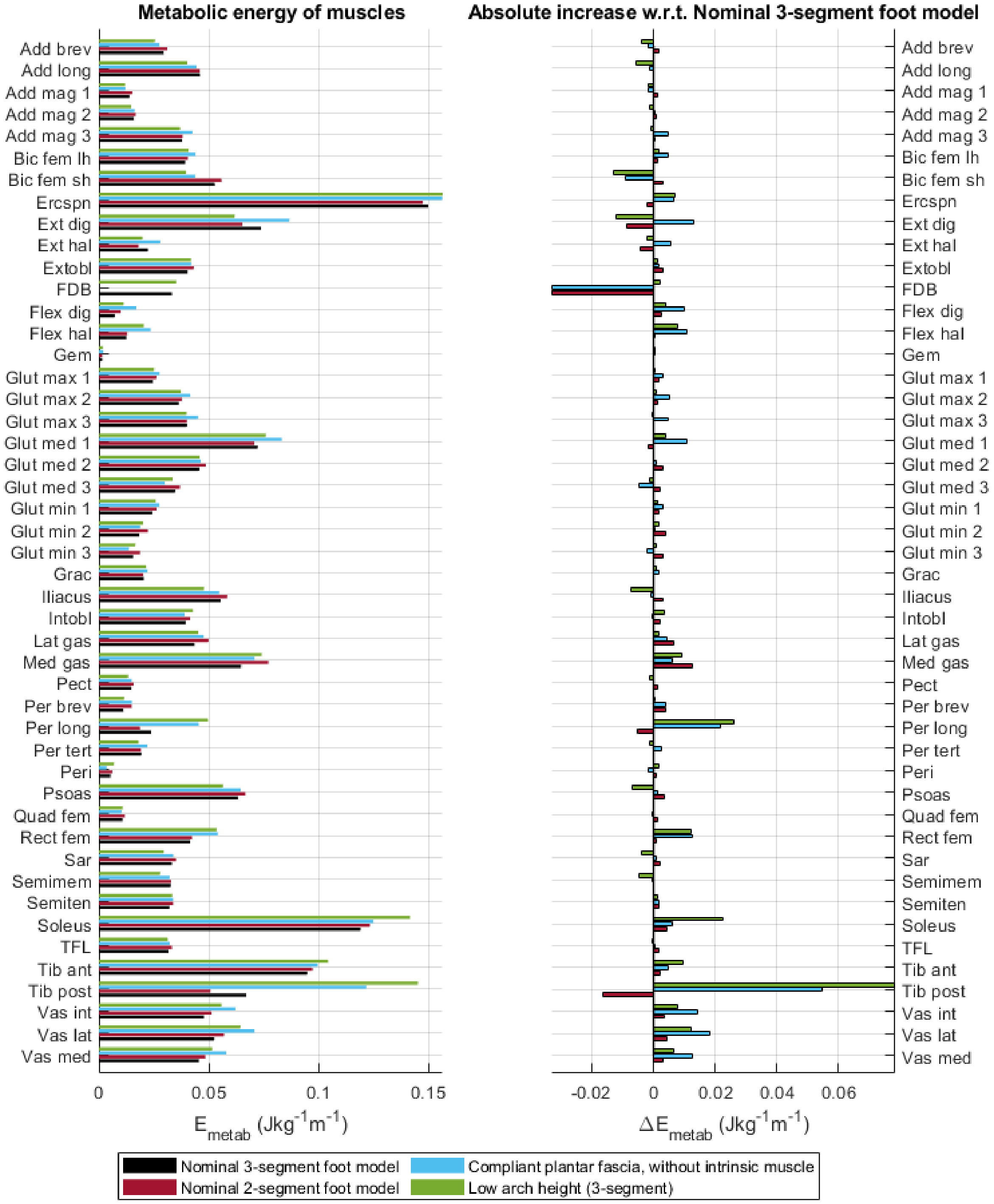


Figure S26 Metabolic energy expenditure of every muscle (right side). Energy is normalised to body mass and distance travelled (cfr. Cost of transport). Left plot shows total values, right graph shows difference with nominal 3-segment foot model.

### S14. Plantar intrinsic muscle nerve block

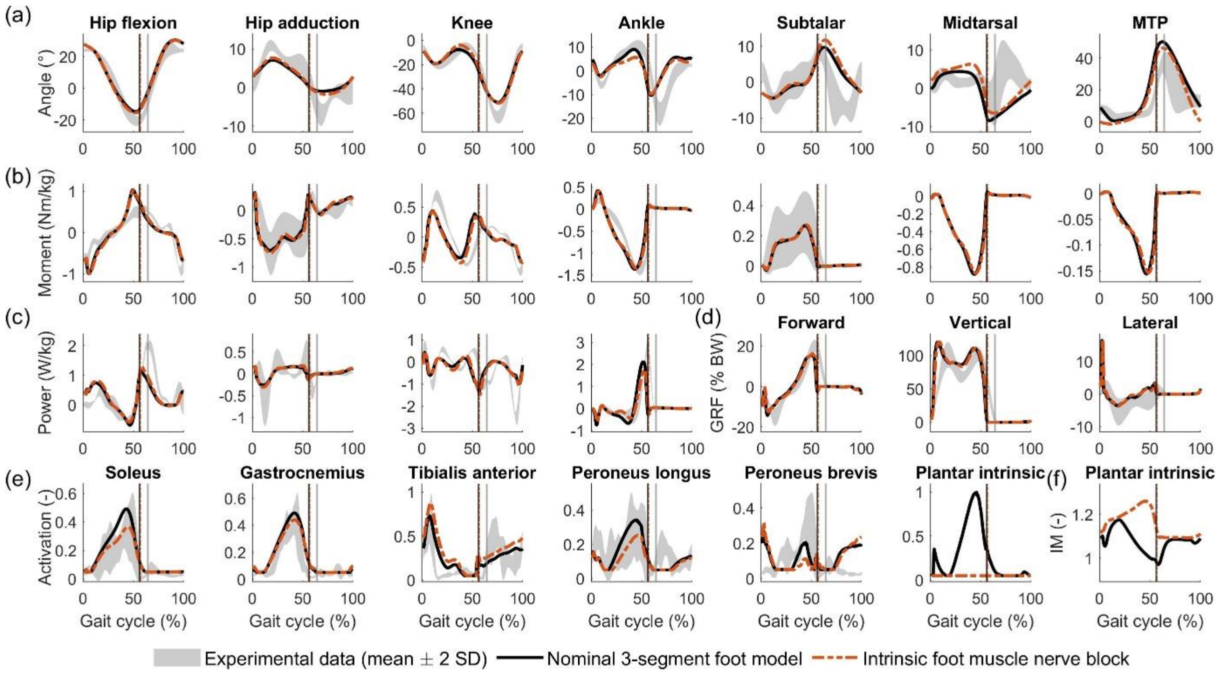


Figure S27 Effect of intrinsic muscle nerve block on gait. (a) Kinematics. (b) Kinetics. (c) Joint powers. (d) Ground reaction forces, expressed as % body weight. (e) Muscle activation. Gastrocnemius indicates the medial gastrocnemius. (f) Fibre length, normalised to optimal fibre length.

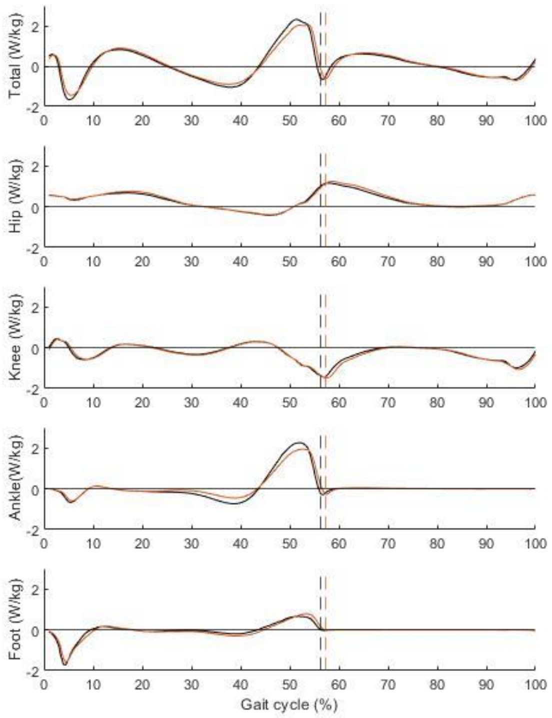


Figure S28 Effects of intrinsic muscle nerve block on joint powers and foot deformation power.

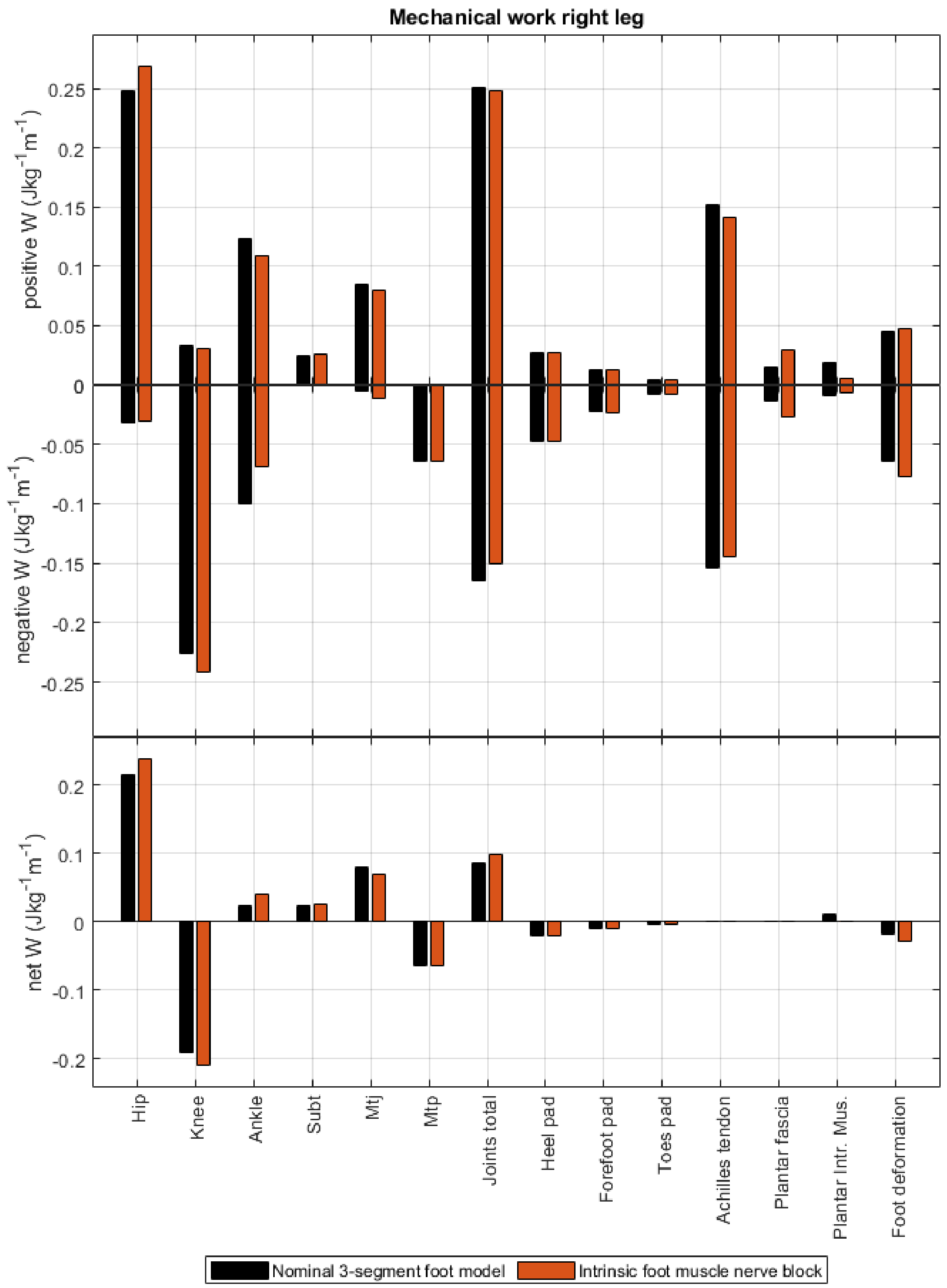


Figure S29 Effect of intrinsic muscle nerve block on positive, negative, and net work around joints and by selected structures.

#### S15. MTP to midtarsal joint energy transfer

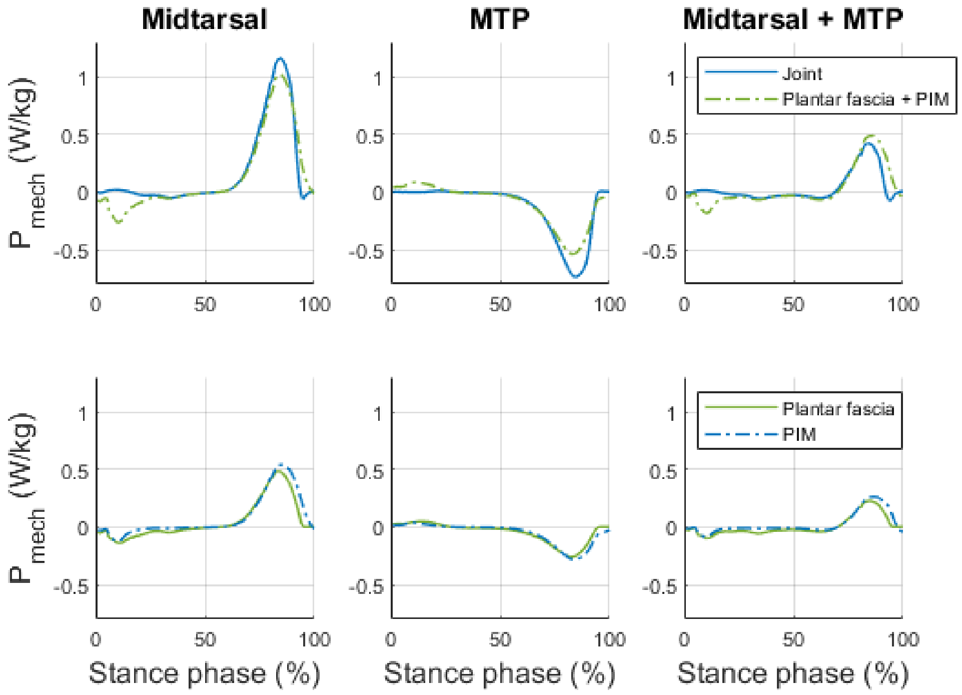


Figure S30 Joint power of midtarsal joint, MTP joint, and summed power of both joints. Plantar fascia and plantar intrinsic muscle (PIM) account for most power around these joints. Separate contributions of plantar fascia and intrinsic muscle have similar magnitude.

#### S16. Contributions of extrinsic foot muscles

The action of the extrinsic foot muscles around the MTP and midtarsal joints was important to obtain plausible joint kinematics with our 3-segment model (Figure S31). Neglecting the muscle actuation of the midtarsal and MTP joints (i.e. passive joints) resulted in exaggerated ankle dorsiflexion and knee flexion throughout stance (Figure S31a). The model with passive foot joints also failed to capture the activity patterns of peroneus longus (Figure S31e). However, in the 2-segment foot model, modelling the action of the extrinsic foot muscles on the MTP joint worsened the prediction of the MTP kinematics compared to the 2-segment foot model with a passive MTP joint. Simulations with a 2-segment foot model with muscle driven MTP joint and extensors resulted in an extended (20°-60°) MTP joint during the entire gait cycle. This is in contrast with the reasonably accurate predictions in MTP angle in the model with the passive MTP joint.

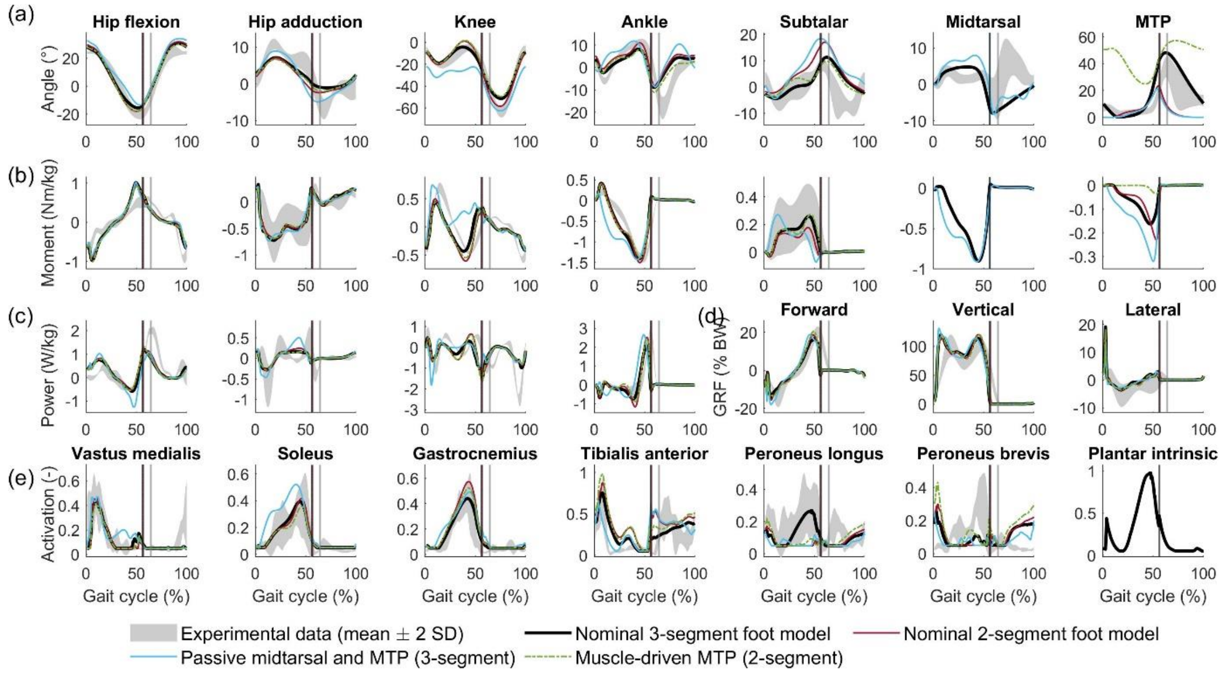


Figure S31: Effect of extrinsic foot muscles. (a) Kinematics. (b) Kinetics. (c) Joint powers. (d) Ground reaction forces, expressed as % body weight. (e) Muscle activation. Gastrocnemius indicates the medial gastrocnemius.

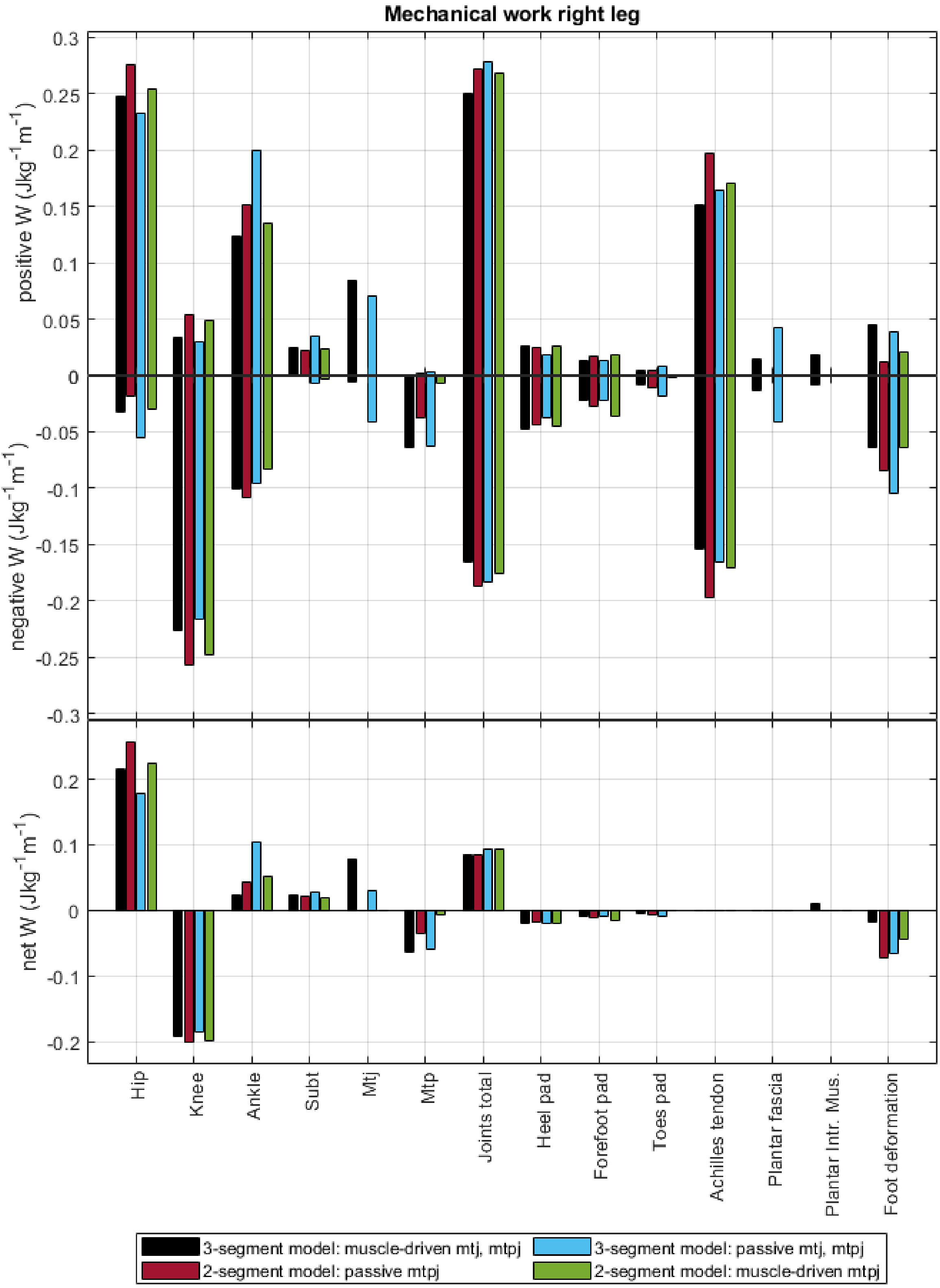


Figure S32 Effect of extrinsic foot muscles on work.
